# Possible Functional Proximity of Various Organisms Based on Taste Receptors Genomics

**DOI:** 10.1101/2022.07.27.501642

**Authors:** Sk. Sarif Hassan, Moumita Sil, Subhajit Chakraborty, Arunava Goswami, Pallab Basu, Debaleena Nawn, Vladimir N. Uversky

**Affiliations:** Department of Mathematics, Pingla Thana Mahavidyalaya, Maligram, Paschim Medinipur, 721140, West Bengal, India; Biological Science Division, Indian Statistical Institute, 203 B.T Road, Kolkata, 700108, West Bengal, India; School of Physics, University of the Witwatersrand, Johannesburg, Braamfontein 2000; Advanced Technology Development Centre, Indian Institute of Technology, Kharagpur, Khargapur, 721302, West Bengal, India; Department of Molecular Medicine, Morsani College of Medicine, University of South Florida, Tampa, FL 33612, USA

**Author notes:** Email addresses:* (Sk. Sarif Hassan), (Moumita Sil), (Subhajit Chakraborty), (Arunava Goswami), (Pallab Basu), (Debaleena Nawn), (Vladimir N. Uversky).

**Keywords:** Taste Receptor, Clusters, Intrinsic Protein Disorder, Shannon variability, Organism proximity

## Abstract

Taste is one of the essential senses in providing the organism a faithful representation of the external world. Taste perception is responsible for basic food and drink appraisal and bestows the organism with valuable discriminatory power. Umami and sweet are “good” tastes that promote consumption of nutritive food, whereas bitter and sour are “bad” tastes that alert the organism to toxins and low pH, promoting rejection of foods containing harmful substances. Not every animal has the same sense of taste as humans. Variation in the taste receptor genes contributes to inter and intra organism differences of taste (sweet/bitter) sensation and preferences. Therefore a deeper understanding was needed to comprehend taste perception by various vertebrates and accordingly elucidate a possible proximity among them. In this study, a total 20 Type-1 (sweet) and 189 Type-2 (bitter) taste receptor complete-amino acid sequences were taken from the 20 vertebrate organisms (18 mammalian, 1 aves, and 1 amphibian). Among 10 primates, 8 including humans were very close based on genomics of taste receptors and rodent organisms viz. the rat and mouse were away from them. This investigation throws light on the similitude and dissimilitude of perception of sweet and bitter taste among 20 different organisms, steered by quantitative analysis of their genomic data. Furthermore, it enlightened that ligand binding affinity of sweet/bitter taste molecules in the taste receptors of any proximal pair of organisms would be similar.

## 1. Introduction

We often come across the phrase “Variety is the spice of life”, physiologically the organ that can perceive this variety of taste is the tongue. A multimodal orchestra of action starting from the tongue leading to neural circuit and ending in the primary gustatory cortex of the brain is responsible for the perception of taste [1, 2]. A taste receptor is a type of cellular receptor which facilitates the sensation of taste [3, 4]. When food or other substances enter the mouth, molecules interact with saliva and are bound to taste receptors in the oral cavity and other locations [5]. Molecules which give a sensation of taste are considered “sapid”. Primarily there are 5 types of taste sensation namely sweet, bitter, sour, salty and umami (the taste of glutamate) [6]. The initial perception of taste begins at the apical end of the taste receptor cells that are found in the taste buds in the mouth [7]. Each bud is garlic shaped and is composed of 50-100 cells [7]. Taste buds are found in structures called papillae on the tongue, palate and on the wall of the throat [8, 9]. Taste buds in the anterior two thirds of the tongue reside in the fungiform papillae, each containing one or a few taste buds. The circumvallate papillae and the foliate papillae reside in the posterior part of the tongue, each of which contains hundreds of taste buds [10]. Humans have about 8-10 thousand taste buds which are replaced every 10 to 14 days. Herbivores like cows have around 25 thousand taste buds, omnivores have 15 thousand and carnivores have the fewest number of test buds. Birds have much fewer taste buds than mammals and fishes have innumerable taste buds both in their mouth and along their lateral lines [11].

Sweet and umami detection help animals find energy-dense nutrients [12]. Bitter detection helps them avoid toxic substances. The question arises whether test perception is similar in all the organisms across the varied taxonomical hierarchy. In this context Dr. Susan Hemsley, senior lecturer in veterinary anatomy and histology at the University of Sydney states that “Different species have different taste buds specialized to detect the things they’re most interested in.” There lie few exceptions in perception of taste. Cats can’t taste sweet things as we humans do [13]. This is an evolutionary trait that everyone has lost in the cat family. One of the genes employed in detecting sweet taste in humans and dogs is inactive in cats, rendering them unable to taste sweet flavors [14]. Artificial sweeteners cannot be perceived by some species of monkey but they can taste natural sugars [15, 16]. Dolphins appear to have no taste receptors other than salty. In a set of revolutionary experiments conducted by R.A. Fischer and his team they introduced phenylthiocarbamide (PTC) into the diet of a few Chimps [17]. PTC is a nauseatingly bitter compound to few humans and others fail to taste it [17]. Three quarters of the Chimps showed displeasure for PTC. Fischer concluded that the bitter taste in both humans and chimps are due to some genetic mutations, which natural selection has maintained in both these species [18]. Kurt Schwenk, a biologist at University of Connecticut states that “The whole story of the evolution of taste is really the evolution of loss of taste” [19]. This variation and loss in certain perception of taste is thought to be an effect of change in lifestyle, a shift in dietary habit [20]. Evolution is a trade of “use it or lose it”, the genes which do not participate in the survival or reproduction are liable to develop mutation leading to malformed protein products. The inability to taste sweetness is profound in other non-feline species belonging to the order Carnivora. In sea-lions, otters and hyenas’ crippling mutations were found in sweet receptors in their respective gene. Also, the mutations were unique in each species. Vampire bats have no sweet receptors and all bats including insectivorous ones lack umami receptors as well [21]. Pandas, a largely vegetarian member of the order Carnivora, lack Umami receptors [22].

As the emphasis has shifted from single gene and protein to the whole genome of organisms, the amount of available data grows in magnitude and thus becomes dependent on mathematical modelling, mathematical analysis and computation. Individual differences in taste, at least in some variants, can be attributed to allelic variants of T1R and T2R genes [23]. Complex stimuli like sugars and most bitter tasting compounds are believed to activate specific G-protein coupled receptors (GPCRs) [24, 25]. The T1R works on a networking basis where TAS1R1+TAS1R3 heterodimer receptor functions as umami receptor sensing L-glutamate, whereas the TAS1R2+TAS1R3 heterodimer receptor functions as a sweet receptor. The TAS2R proteins function as bitter taste receptors [26, 27].

In this present study, we attempted to illuminate the nearness of Type-1 (sweet) and Type-2 (bitter) taste receptors across different vertebrate organisms by extracting various quantitative features of the taste receptor sequences. The yardstick on which the nearness of the organisms would be assessed includes the multiple sequence alignment, the analyses of the amino acid frequency, polarity of amino acids, physicochemical properties, biophysical properties, and single nucleotide polymorphisms. The resemblance and/or divergence of the perception of sweet and bitter taste among 20 different species had been explored in this study, in reference to genomic data. Hands-on scientific research and experimentation is additionally required for validating the proposed hypothesis.

## 2. Data and methods

### 2.1. Data acquisition

A set of 20 Type-1 (sweet) and 189 Type-2 (bitter) taste receptor complete-amino acid sequences was assembled from the following organisms (Table-1) (UniProt) [28].

**Table 1:**
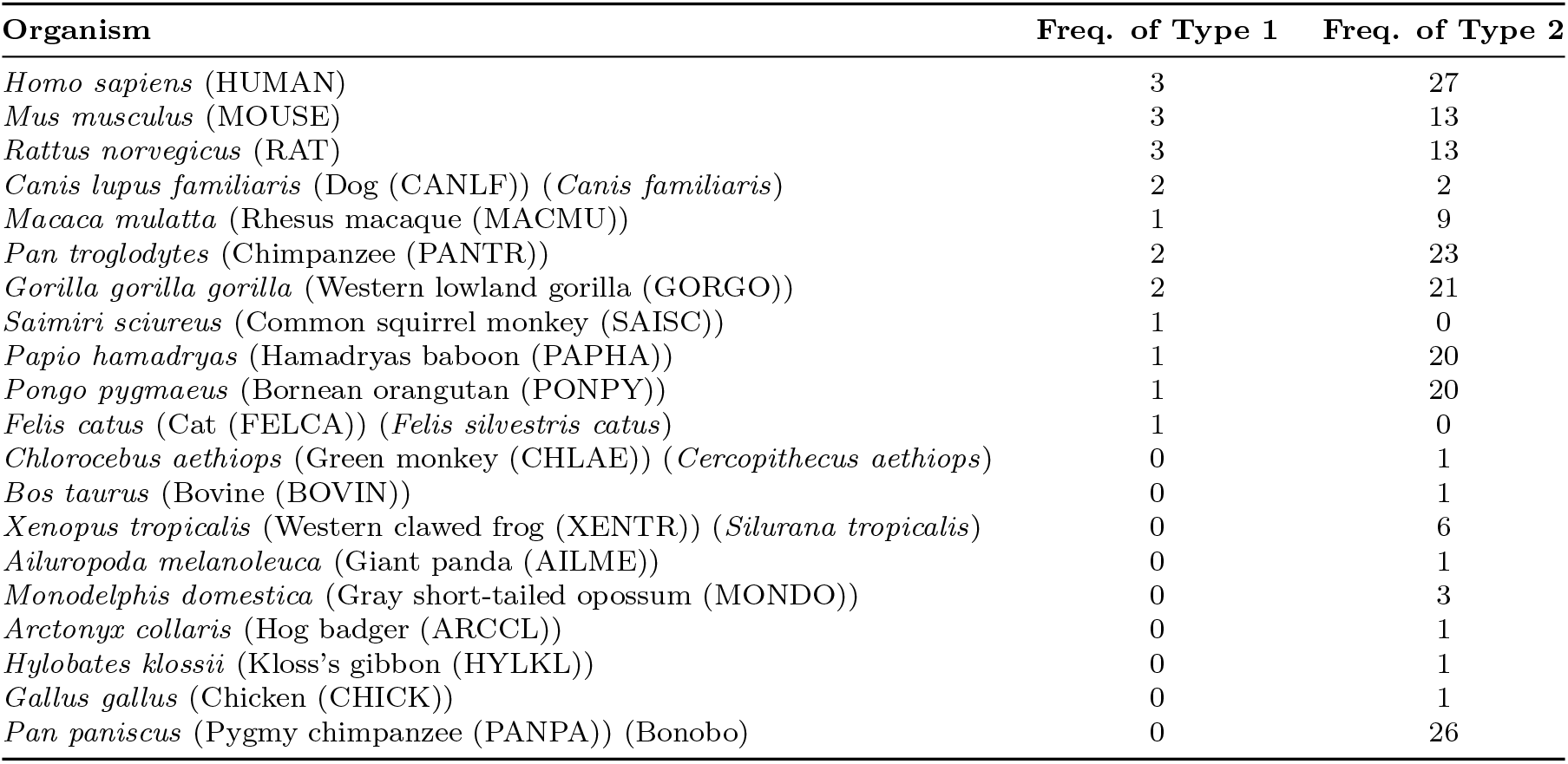
List of 20 organisms from which Type-1 and Type-2 taste receptor sequences with respective frequencies (Freq.) were considered.

List of all the accession numbers of all taste receptor sequences was made available as a **Supplementary file-1**. Furthermore, it was noted that the following pairs of sequences were 100% identical:

(sp—Q646H0—TA2R1_PANTR, sp—Q646G9—TA2R1_PANPA), (sp—Q646B5—T2R10_PANTR, sp—Q646D7—T2R10_PANPA), (sp—Q646D2—TA2R3_PANPA, sp—Q645Y5—TA2R3_GORGO), (sp—Q646D2—TA2R3_PANPA, sp—Q646A7—TA2R3_PANTR), (sp—Q645Y5—TA2R3_GORGO, sp—Q646A7—TA2R3_PANTR), (sp—Q646A9—T2R39_PANTR, sp—Q646C8—T2R39_PANPA), (sp—Q645T6—T2R50._MACMU, sp—Q646G2—T2R50_PAPHA), and (sp—Q646E4—T2R50_PANPA, sp—Q646C3—T2R50_PANTR).

### 2.2. Methods

#### 2.2.1. Quantitative features extracted from Taste receptors

##### Deviation from randomness

A statistical deviation from randomness in the spatial arrangement of amino acids is studied by analyzing the clustering properties [29, 30]. First, every amino acid in a given taste receptor amino acid sequence was identified as polar(P) or non-polar(Q) [29]. Thus, every taste receptor amino acid sequence became a binary sequence with two symbols P and Q. We test the null hypothesis that such a string of P and Q is a random string with no-correlation between the placements of polar and non-polar amino acids. To test our null hypothesis we perform the following bootstrap analysis in three steps:

1. First we note the average cluster size (*m*) of a binary sequence.
2. We randomly permutated each binary sequence 5000 times and noted the mean (*μ*) and variance (*σ*^2^) of average cluster sizes. We define: 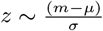 and we note the value of ‘*z*’.
3. We perform the above two steps for all the binary sequences. We check our null hypothesis by performing a student-t test to check if mean of *z* is significantly different from 0.

##### Evaluating intrinsic protein disorder predisposition

Per-residue disorder propensity within the amino acid sequences of two types of taste receptors was evaluated by PONDR@ VSL2, which is one of the more accurate standalone disorder predictors [31, 32]. The per-residue disorder predisposition scores are on a scale from 0 to 1, where values of 0 indicate fully ordered residues, and values of 1 indicate fully disordered residues. Values above the threshold of 0.5 are considered disordered residues, whereas residues with disorder scores between 0.25 and 0.5 are considered highly flexible, and residues with disorder scores between 0.1 and 0.25 are considered as moderately flexible.

##### Amino acid frequency based Shannon entropy and Shannon variability

*Shannon entropy:* How conserved/disordered the amino acids are organized over a protein sequence is addressed by the information-theoretic measure known as Shannon entropy (SE). For a given amino acid sequence of length *l*, the conservation of amino acids is calculated as follows:

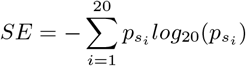

where 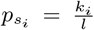 represents the frequency distribution of each amino acid (AAC) present in a taste receptor sequence was determined using standard MATLAB bioinformatics routine. [33, 34].

*Shannon Variability:* Shannon entropy is deployed to estimate variability of amino acid residues at each residue position across all the taste receptor sequences [35]. For a multiple protein sequence alignment, the Shannon variability (H) for every position is defined as follow:

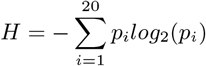

where *p_i_* represents the fraction of residues of amino acid type *i. H* ranges from 0 (only one residue present at that position) to 4.322 (all 20 residues are equally represented in that position). Typically, positions with *H* > 2.0 are considered variable, whereas those with *H* < 2 are considered conserved. Highly conserved positions are those with *H* < 1.0 [35, 36].

##### Determining SNPs, Pathogenicity, Stability

Without loss of generality, reference sequence Taste receptor Type-1 member 1 (Q7RTX1—TS1R1_HUMAN) and Taste receptor Type-2 member 21 (sp—Q8MJU6—TA2R1_CHLAE) were taken to assess the single nucleotide polymorphism (SNP) across Type-1 and Type-2 taste receptors respectively. The similar amino acid residual non-synonymous changes based on multiple sequence alignment executed by the Clustal Omega web-server with reference to the reference taste receptor sequences was termed here as ‘SNPs’ [37, 38].

Mutations in proteins are linked with the development of various genetic diseases. Computational tools for the prediction of the effects of mutations on protein function are very important for analysis of single nucleotide variants and their prioritization for experimental characterization. The PREDICTSNP web-server uses the six best performing tools MAPP, nsSNPAnalyzer, PANTHER, PhD-SNP, PolyPhen-1, PolyPhen-2, SIFT, and SNAP to predict pathogenicity for a given mutation. The consensus predictions for mutations were found to be accurate and robust [39].

Protein function is directly governed by associated protein structure and a single mutation on the amino acid residue may cause a severe change in the protein structure and thus, lead to disruption of function. Therefore, predicting the protein stability changes can provide several possible candidates for the novel protein designing. iStable takes either in the form of structure or sequence as input [40]. In the present study, taste receptor protein sequences were taken as input and found the stability for each SNPs.

##### Structural and physicochemical features

Structural and physicochemical descriptors (PC) of a protein sequences have been widely used to characterize sequences and predict structural, functional, expression, and interaction profiles of proteins. iFeature, is an widely used Python-based web-server for generating various numerical features for protein sequences [41]. A list of 1182 features which were deployed in the present study was summarized in the following Table 2 [41].

**Table 2:** List of features of dimension 1182 calculated by iFeature for all the taste receptor sequences.

| List of various features (1182 dimensional) calculated by iFeature |  |  |
| --- | --- | --- |
| Descriptor groups | Descriptor | Dimension |
| Amino acid composition | Amino acid composition (AAC) | 20 |
|  | Dipeptide composition (DPC) | 400 |
| Grouped amino acid composition | Grouped Amino Acid Composition (GAAC) | 5 |
|  | Grouped Dipeptide Composition (GDC) | 25 |
| Autocorrelation | Moran Autocorrelation | 240 |
| C/T/D | C/T/D Composition (CTDC) | 39 |
| Conjoint Triad | Conjoint Triad | 343 |
| Quasi-sequence-order | Sequence-Order-Coupling Number (SoC number) | 60 |
| Pseudo-amino acid composition | Pseudo-Amino Acid Composition (PAAC) | 50 |

##### Biophysical features

Nine biophysical features (BP) viz. amino acid frequency based Shannon entropy, molecular weight, theoretical pI, total number of negatively charged residues (Asp + Glu), total number of positively charged residues (Arg + Lys), extinction coefficient, instability index, aliphatic index, and grand average of hydropathicity (GRAVY) of all taste receptor sequences were calculated using the web-server ‘ProtParam’ [42, 43].

#### 2.2.2. Statistical significance and correlation analysis

A well-known non-parametric test to compare two independent groups is the *Mann Whitney U test,* when the distribution is not normal [44, 45, 46]. In this study, two groups were considered, and they were Type-1 and Type-2 taste receptor sequences. Mann Whitney U test was carried out to check whether features such as structural and physicochemical properties, biophysical properties, and so on are significantly different between two types of taste receptor sequences. It was noted that significant differences between the number of Type-1 (20) and Type-2 (189) taste receptor sequences may produce unreliable results from statistical analysis. In order to avoid that, a combination of 20 Type-2 taste receptor sequences were randomly chosen to form a subset and 300 such subsets from the set of 189 Type-2 sequences were generated. A feature was concluded to have significant differences (for one-tailed test at significance level of 0.01), if differences were found in at least 90% of the total number of comparisons between Type-1 and Type-2.

In finding correlation between all possible pair of significant features (for each Type), Pearson correlation coefficients were computed.

#### 2.2.3. Hierarchical relationship of taste receptor sequences

- *Hierarchical relationship based on multiple sequence homology:* Multiple sequence alignment (MSA) among the taste receptor sequences was executed by Clustal Omega and further reverified by Uniprot-Align web-servers and consequently, nearest neighbour phylogenetic trees were developed [47, 48].
- *Hierarchical relationship based on Polarity:* The sequence homology of polar-nonpolar (PNP) sequences for every taste receptor sequence was derived from Clustal Omega web-server and then nearest neighbour hierarchical relationships among sweet and bitter taste receptor sequences were deduced.
- *Hierarchical relationship based on amino acid frequency (AAC), physicochemical, and biophysical features:* For each taste receptor sequence, a twenty-dimensional frequency-vector considering the frequency of standard twenty amino acids can be obtained. The distance (Euclidean metric) between any two frequency vectors was calculated. By having a distance matrix (Euclidean distance), a hierarchical relationship was developed based on the nearest neighbour-joining method using the standard routine in *Matlab-2022a* [49]. Likewise, distance matrices for the normalized (standard) physicochemical and biophysical features were derived and accordingly hierarchical relationships among the taste receptor sequences were drawn.

#### 2.2.4. Clustering of organisms using UPGMA

UPGMA (unweighed pair group method with arithmetic mean), an agglomerative hierarchical clustering method was deployed to cluster (circular phylogram) various organisms, based on distance matrix formed by several features from taste receptor sequences [50, 51]. Following two definitions were given to clarify the clustering of organisms.

*Disjoint cluster*: cluster containing only 2 organisms.

*Extended cluster*: cluster containing more than 2 organisms.

- *Distance matrix formation for SNPs:* Number of different mutations (ND) was found between two organisms *i* and *j* (*i,j* = 1, 2,… 11 for Type-1 and *i,j* = 1, 2,… 18 for Type-2 and *i* ≠ *j*). ND was obtained by subtracting common mutations (intersection) from union of all mutations of *i* and *j*.
- *Distance matrix formation for PNP and MSA:* Median similarity values of all possible pairwise combinations of sequences between two organisms were considered to form species wise similarity matrices from sequence wise similarity matrices. Distances were obtained by subtracting similarity values from 100.
- *Distance matrix formation for AAC, BP, and PC:* Each feature of AAC, PC, and BP were normalized separately. Sequences of the same organisms were considered as one group. Euclidean distances between all possible pairwise combinations of sequences between two groups (same organism) *i* and *j* (*i,j* = 1, 2,… 11 for Type-1 and *i,j* = 1, 2,… 18 for Type-2) were calculated and their median was finally considered as distance between group *i* and *j*.

*Cumulative clustering of organisms for Type-1 and Type-2 features:* Each distance matrix was normalized by dividing with their maximum values and five matrices (BP, PC, PNP, MSA and SNP) were added to obtain cumulative clustering of Type-1 and Type-2.

*Clustering of common organisms of Type-1 and Type-2:* Nine organisms viz. HUMAN, MACMU, PAPHA, CANLF, PONPY, MOUSE, and RAT had both Type-1 and Type-2 taste receptors. Cumulative distance matrices of Type-1 and Type-2 for these nine organisms were added to derive clustering of common organisms.

*Possible functional proximity of organisms:* Proximal nearness among all the 20 organisms was obtained by taking union of the cumulative clustering for Type-1 and Type-2.

## 3. Results and Discussion

### 3.1. Deviation from randomness leads to the organized polarity

The *p*–value obtained for Type-1 and Type-2 were *p* = 2.2*e* — 6 (with statistic=6.72) and *p* = 1*e* — 4 (with statistic=3.80), respectively. Hence, in both cases the null hypothesis was rejected. The strength of the effect is the following: for Type-1 polar-nonpolar sequences *Mean*(*z*) = 1.31 and Type-2 polar-nonpolar sequences *Mean*(*z*) = 0.28 (**Supplementary file-2**). This showed a higher deviation from randomness for Type-1 polar-nonpolar sequences than that of Type-2 sequences. Mann-Whitney-U test was deployed to check the difference in *z*-values between Type-1 and Type-2 sequences, and a significant difference was observed that a significant difference with *p* = 6.61*e* — 06, statistic=769. The following histograms showed the difference between the standard normal and the real distribution of *z* (Figure 1).

**Figure 1:**
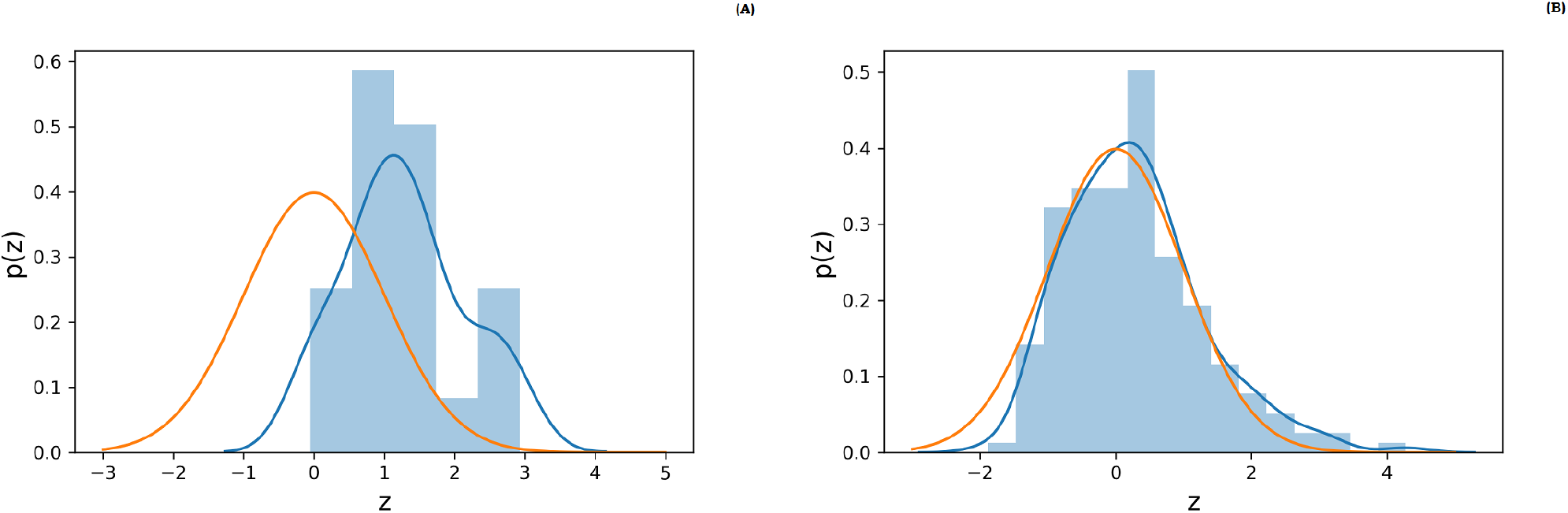
Difference between the standard normal and the distribution of average cluster sizes of the binary versions of Type-1 (A) and Type-2 (B) taste receptor sequences. Orange: standard normal, Blue: the histogram and kernel density of mean cluster sizes.

Furthermore, organism level deviation in randomness by limiting our analysis to sequences belonging to a particular organism was carried out. It was noticed that for all Type-2 taste receptor polar-nonpolar sequences *Mean*(*z*) = 0.224 for HUMAN, *Mean*(*z*) = 0.38 for PONPY, and *Mean*(*z*) = 0.44 for MOUSE, and *Mean*(*z*) = 0.613 for RAT. This observation showed an organism-wise departure from randomness for Type-2 sequences, which was higher in rodent organisms. It turned out that polarity (polar-nonpolar) sequences of Type-1 and Type-2 taste receptors of all the 20 organisms were deviated from random binary sequences. In other words, an organized polarity of taste receptor sequences of various organisms was noticed.

### 3.2. Protein intrinsic disorder predisposition of taste receptors

Protein intrinsic disorder predisposition analysis revealed that the propensity of Type-1 and Type-2 receptors from different organisms for intrinsic disorder can vary significantly, especially in the C- and N-termini of the Type-2 receptors (Figure 2A) and 2B). Although the Type-1 receptors were predicted a bit more disordered than the Type-2 receptors (Figure 2C), their disorder profiles were characterized by lower variability. Importantly, for both receptor types, the largest variability was observed within the disordered or flexible regions of these proteins (i.e., regions characterized by the predicted disorder scores exceeding the 0.5 threshold and regions with disorder scores between 0.15 and 0.5) (Figure 2A) and 2B). This is an important observation suggesting that natural variability of Type-1 and Type-2 taste receptors proteins is shaping their structural flexibility.

**Figure 2:**
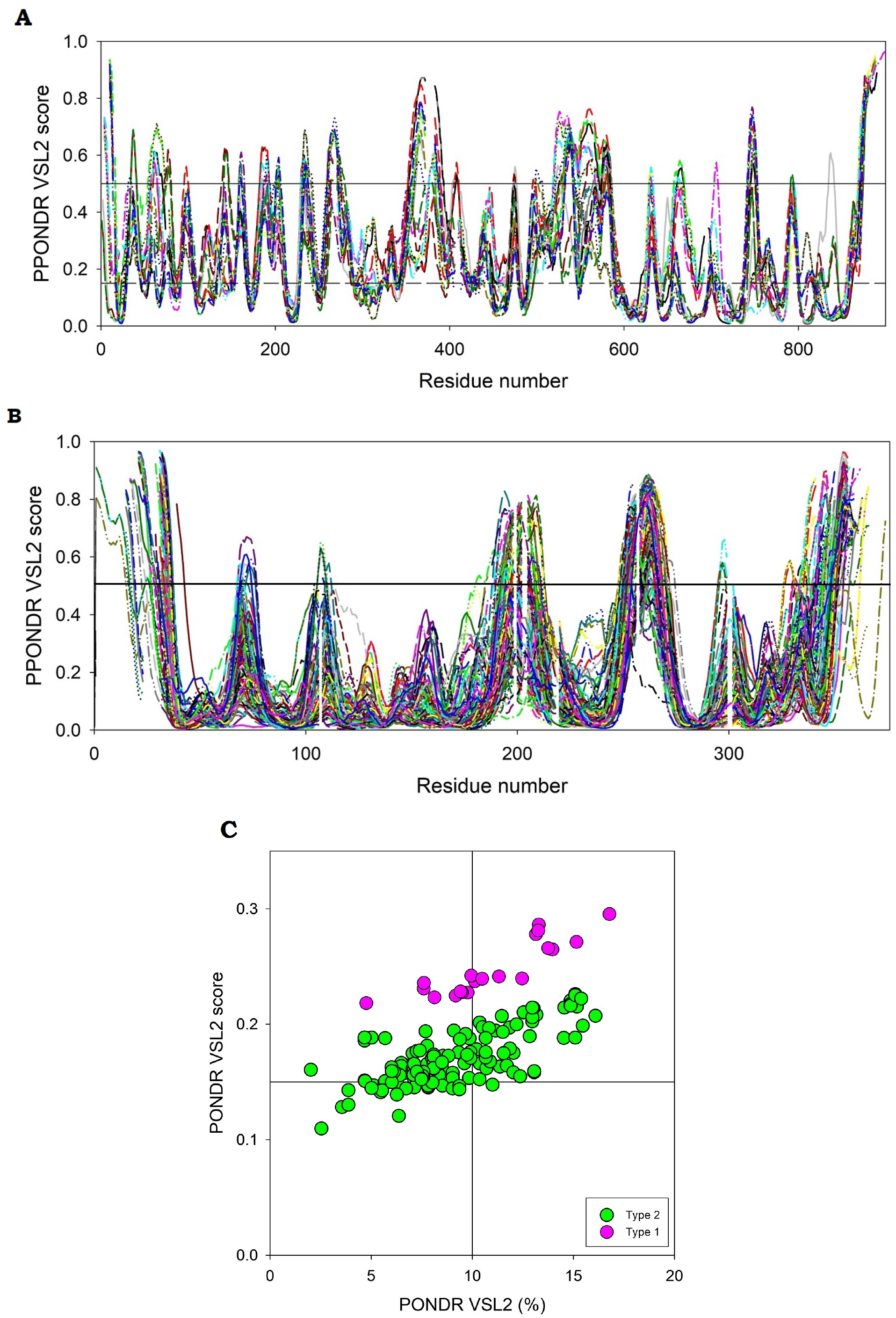
Per-residue intrinsic disorder profiles generated for Type-1 (A) and Type-2 (B) taste receptors from twenty species by PONDR@ VSL2, and global intrinsic disorder predisposition analysis of 20 Type-1 (sweet, pink circles) and 189 Type-2 (bitter, green circles) taste receptors based on the average disorder score and percent of predicted disordered residues as evaluated by PONDR@ VSL2 (C).

### 3.3. Shannon variability of taste receptors

Variability of each amino acid residues at each position was calculated based on Shannon entropy (Figure 3).

**Figure 3:**
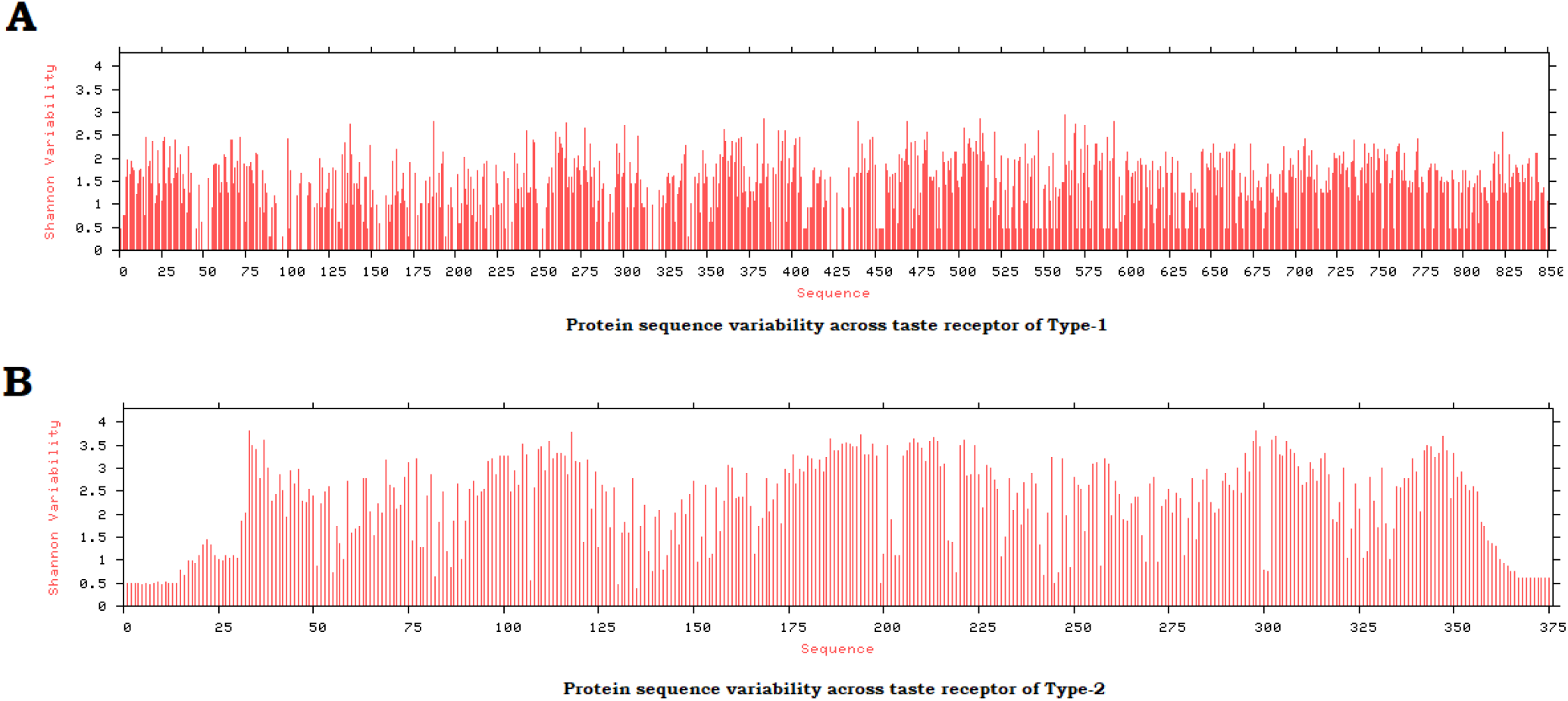
Protein sequence variability across all taste receptor sequences (Type-1: (A) and Type-2: (B))

It was noticed that 17.53% of the amino acid residues were found varied across the 20 taste receptor sequences of Type-1, whereas 63.47% amino acid residues were varied over 189 Type-2 taste receptors (Table 3). These observations are in line with the results of intrinsic disorder predisposition analysis, according to which the Type-2 receptors were more variable than the type-1 receptors (cf. data shown in Figure 2A) and 2B).

**Table 3:** Shannon variability of amino acid residue conservation across the Type-1 and Type-2 taste receptors.

| Shannon Variability (%) |  |  |  |
| --- | --- | --- | --- |
| Types | Highly conserved | Conserved | Variable |
| **Type-1** | 27.53 | 54.94 | 17.53 |
| **Type-2** | 14.4 | 22.13 | 63.47 |

### 3.4. Mutations lead to clustering of organisms

Based on the amino acid sequence homology across all the taste receptors of each type, similar residual changes were accounted for as SNPs. A total of 294 and 22 SNPs were detected in Type-1 and Type-2 taste receptor sequences, respectively (**Supplementary file-3**). Large variation between the total number of SNPs in two types of taste receptors clearly showed wide sequence diversity of Type-2 taste receptors, whereas Type-1 taste receptors were found to be more conserved as compared to Type-2 taste receptors. In Type-1, among 294 SNPs, 248 of them were neutral and in most cases (192 SNPs) structural stability predicted to be decreased, whereas 11 deleterious mutations were predicted to increase the structural stability. In case of Type-2, only two mutations viz. L197M and A227V were turned out to be neutral.

In obtaining the clusters based on the number of disagreement of SNPs between two organisms, lists of all mutations detected in each organism (for both the types) were noted, and consequently, clustering of organisms was made.

For both types of taste receptors, RAT and MOUSE belonged to one single cluster. HUMAN, GORGO and PANTR belonged to an extended cluster (EC_1_HGP) in Type-1 with very small distances among them (Figure 4). Furthermore, PONPY and PAPHA along with MACMU formed an extended cluster (EC_1_PoPaM) which was away from EC_1_HGP. Distances among PONPY, PAPHA, and MACMU were very small. In contrast, GORGO, PANTR, PONPY, and PAPHA made an extended cluster in Type-2 with negligible distances among them. HUMAN was immediately next to the cluster consisting of GORGO, PANTR, PONPY, and PAPHA in Type-2. Unlike Type-1, MACMU was significantly distant from PONPY and PAPHA in Type-2, as noticed in Figure 4B. CANLF was adjacent to EC_1_HGP in Type-1, while CANLF was away from HUMAN, GORGO and PANTR in Type-2.

**Figure 4:**
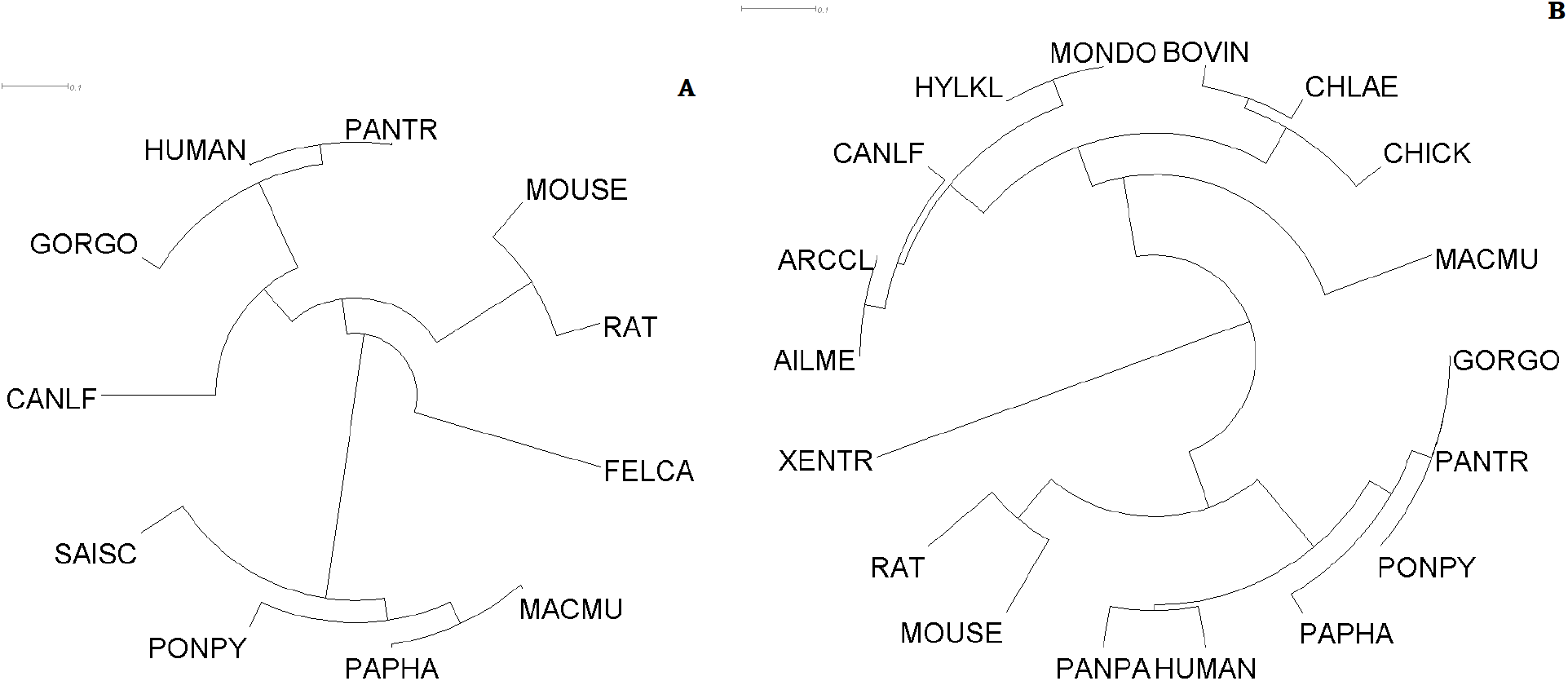
Clustering of organisms based on disagreement of SNPs of Type-1 (A) and Type-2 (B) taste receptors

Also, it was noticed that no SNPs were detected at 22 positions in the T2R19_HUMAN with reference to the TA2R1_CHLAE sequence (Figure 5).

**Figure 5:**
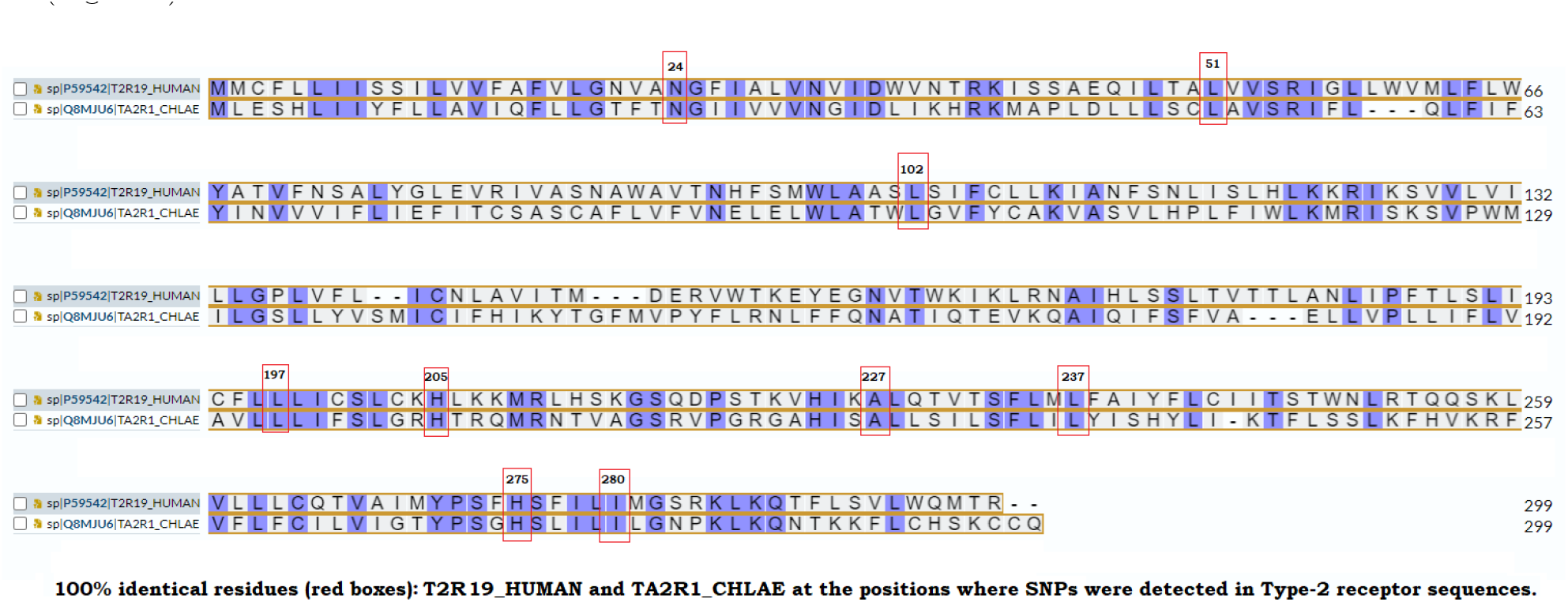
Amino acid sequence homology between T2R19_HUMAN and TA2R1_CHLAE Type-2 taste receptor sequences

### 3.5. Amino acid sequence based homology, hierarchical relationships, and clustering

Based on the amino acid sequence homology (MSA), three members (TS1R1, TS1R2, and TS1R3) of Type-1 from different organisms formed three disjoint clusters as observed in (Figure 6). Note that, the hierarchical relationship among the Type-2 taste receptors is shown in the **Supplementary file-4**.

**Figure 6:**
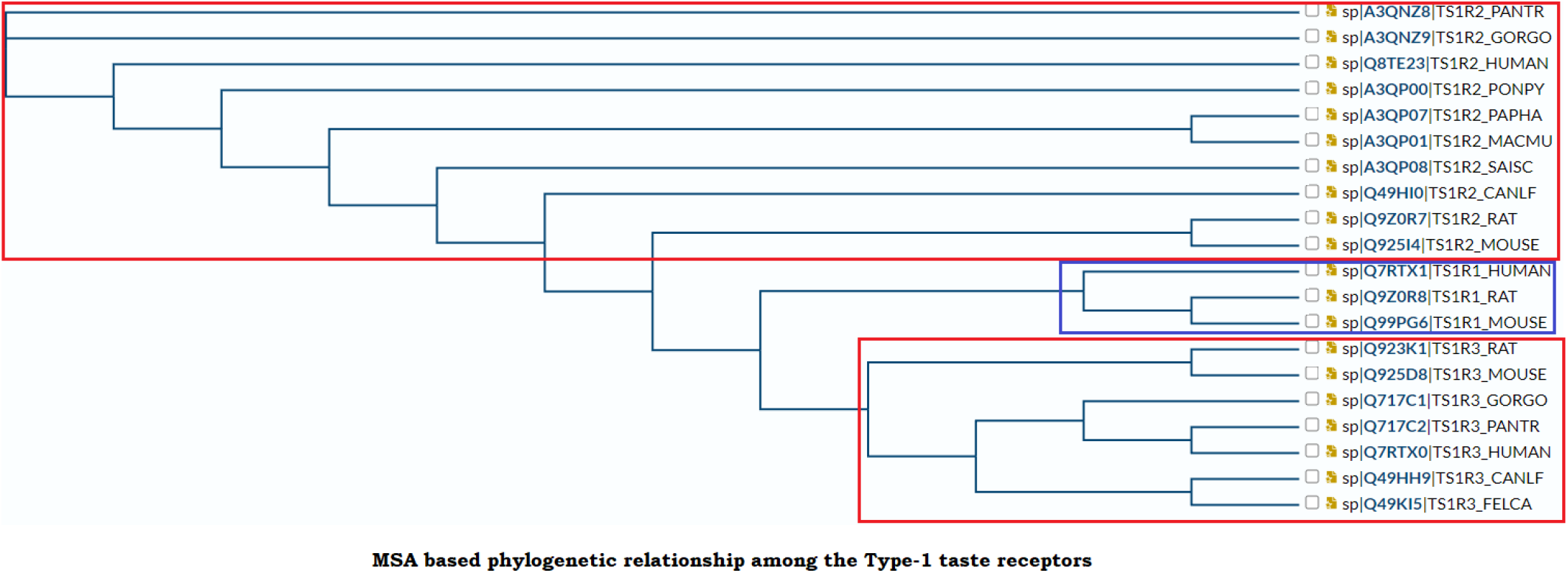
Hierarchical relationship of Type-1 taste receptors based on amino acid sequence homology (multiple sequence alignment)

In Type-1, GORGO and PANTR formed a distinct cluster (EC_1_GP) (Figure 7). Similar to SNP Type-1 (Figure 4A), distances among PONPY, PAPHA, and MACMU (components of EC_1_PoPaM) were very small. HUMAN was quite far from PANTR as well as EC_1_PoPaM. On the other hand, all the above-mentioned organisms viz. GORGO, PANTR, PONPY, PAPHA, MACMU, and HUMAN being almost equidistant to each other, formed an extended cluster in Type-2 (EC_2_GPPoPaMH). CANLF was closer to all above-mentioned organisms in Type-1 than Type-2 (Figure 7).

**Figure 7:**
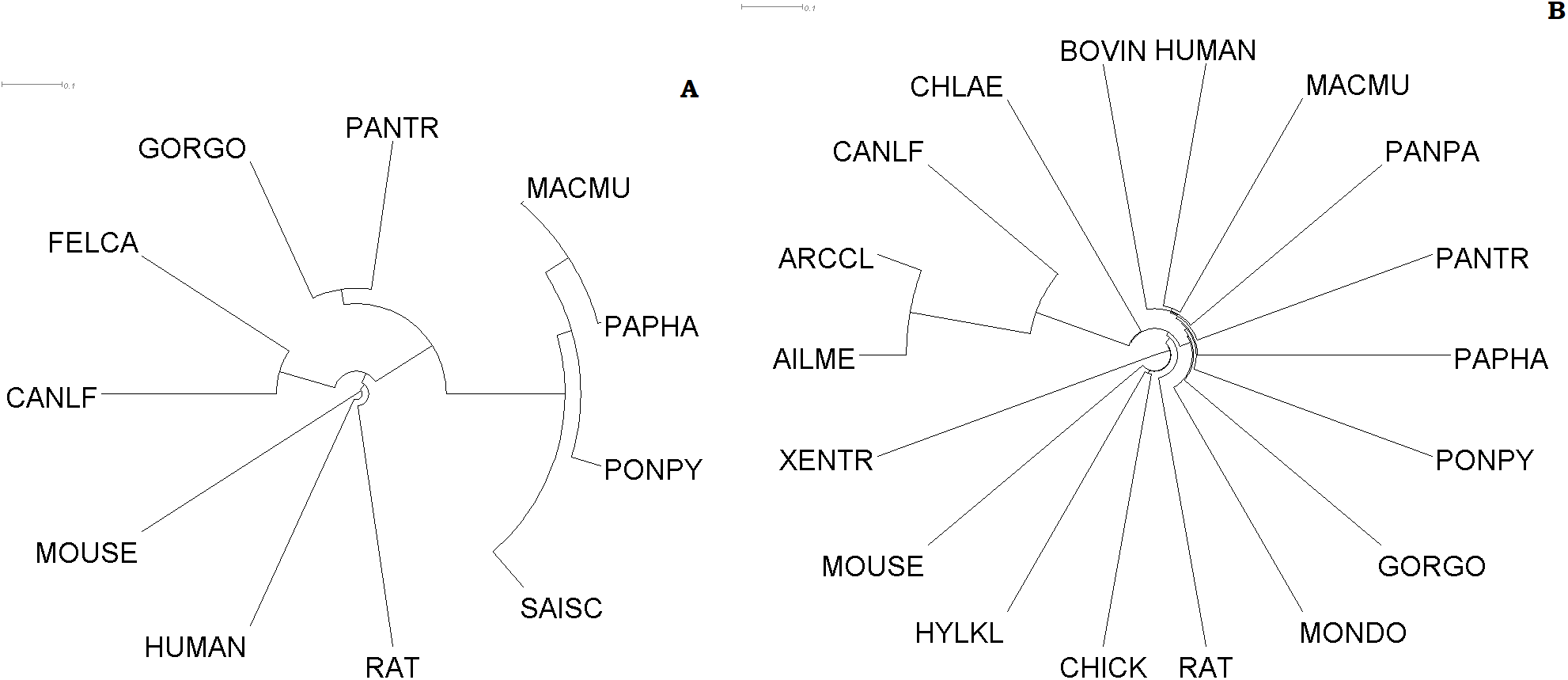
Clustering of organisms based on amino acid sequence homology of Type-1 (A) and Type-2 (B) taste receptors

### 3.6. Polarity based homology, hierarchical relationships, and clustering

It was noticed that all the 20 polar-nonpolar (PNP) binary sequences of Type-1 were distinct. The lowest percentage (72.16%) of similarity was noticed for the pair (sp—Q717C1—TS1R3_GORGO, sp—A3QP00—TS1R2_PONPY) and the highest percentage (99.65%) was noticed for sp—Q7RTX0—TS1R3_HUMAN and sp—Q717C2—TS1R3_PANTR. Furthermore, it was found that among 189 Type-2 sequences, 17 groups of 100% identical sequences were found (Table 4). It was observed from Table 4 that polar-nonpolar binary sequences (Type-2) of PANTR and PANPA were 100% identical in 14 groups. Only in one group (Group-15) polar-nonpolar sequences of MACMU and PAPHA were found to be identical. For Type-2, the lowest percentage (58.04%) of similarity was noticed for the pair sp—Q646G3—T2R42_PAPHA and tr—Q2AB62—Q2AB62_XENTR.

**Table 4:** List of 17 groups of Type-2 taste receptor sequences having identical polar-nonpolar amino acid residues.

| List of Type-2 taste receptor sequences having identical polar-nonpolar amino acid residues |  |  |
| --- | --- | --- |
| **Group-1**<br>sp—Q9NYW7—TA2R1_HUMAN<br>sp—Q646H0—TA2R1_PANTR<br>sp—Q646G9—TA2R1_PANPA | **Group-6**<br>sp—Q646C1—T2R30_PANTR<br>sp—Q646E2—T2R30_PANPA | **Group-12**<br>sp—P59537—T2R43_HUMAN<br>sp—Q646B4—T2R43_PANTR |
| **Group-2**<br>sp—Q646B5—T2R10_PANTR<br>sp—Q646D7—T2R10_PANPA | **Group-7**<br>sp—Q646A9—T2R39_PANTR<br>sp—Q646C8—T2R39_PANPA | **Group-13**<br>sp—Q646D5—TA2R5_PANPA<br>sp—Q646A4—TA2R5_PANTR |
| **Group-3**<br>sp—Q646B6—T2R13_PANTR<br>sp—Q646D8—T2R13_PANPA | **Group-8**<br>sp—Q9NYW5—TA2R4_HUMAN<br>sp—Q646D3—TA2R4_PANPA<br>sp—Q646B0—TA2R4_PANTR | **Group-14**<br>sp—P59544—T2R50_HUMAN<br>sp—Q646A1—T2R50_GORGO |
| **Group-4**<br>sp—Q9NYV7—T2R16_HUMAN<br>sp—Q646D1—T2R16_PANPA<br>sp—Q646B3—T2R16_PANTR<br>sp—Q645Y4—T2R16_GORGO | **Group-9**<br>sp—Q646B1—T2R40_PANTR<br>sp—Q646D4—T2R40_PANPA | **Group-15**<br>sp—Q645T6—T2R50_MACMU<br>sp—Q646G2—T2R50_PAPHA |
| **Group-5**<br>sp—Q646E3—T2R20_PANPA<br>sp—Q646C2—T2R20_PANTR | **Group-10**<br>sp—Q646B2—T2R41_PANTR<br>sp—Q646C7—T2R41_PANPA | **Group-16**<br>sp—Q646A5—T2R60_PANTR<br>sp—Q646D0—T2R60_PANPA |
|  | **Group-11**<br>sp—Q646B8—T2R42_PANTR<br>sp—Q5Y4Z2—T2R42_PANPA | **Group-17**<br>sp—Q646D2—TA2R3_PANPA<br>sp—Q645Y5—TA2R3_GORGO<br>sp—Q646A7—TA2R3_PANTR |

Similar to Figure 6, here also three members of Type-1 formed three separate clusters as marked in Figure 8. Hierarchical relationship of Type-2 taste receptors based on PNP based alignment is supplied in the **Supplementary file-5**.

**Figure 8:**
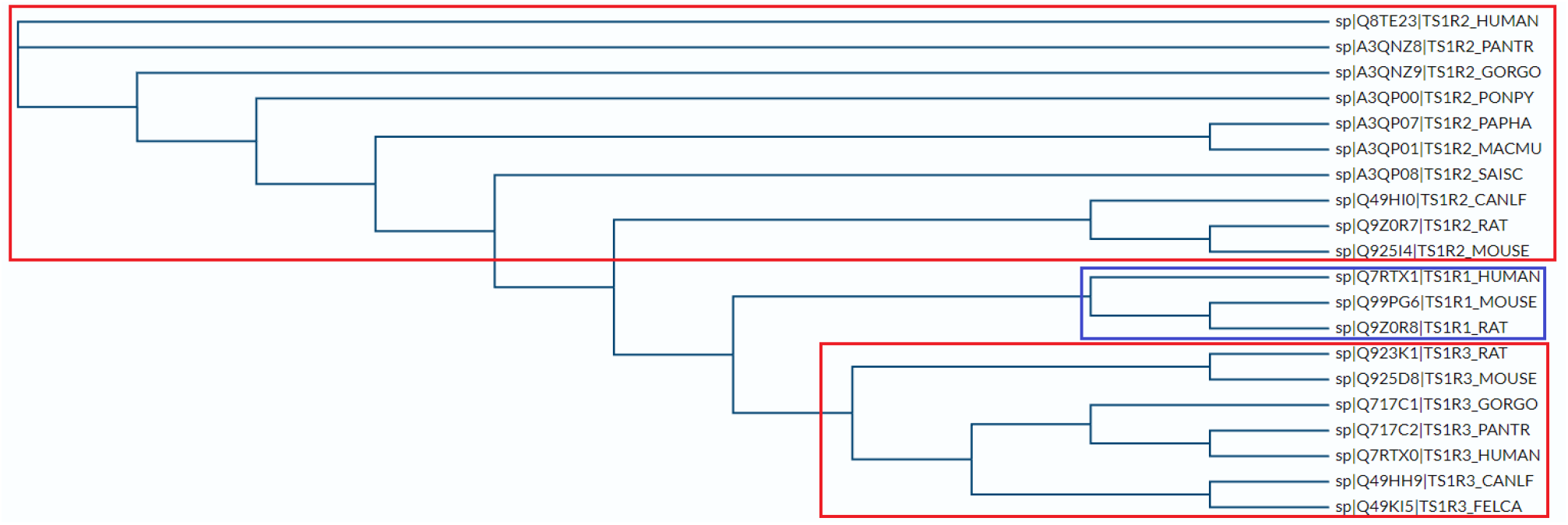
Hierarchical relationship of Type-1 taste receptors based on polar-nonpolar residue based alignment

From Figure 9A it was noticed that in Type-1, GORGO and HUMAN formed a distinct cluster. HUMAN was far apart from EC_1_PoPaM. Although EC_1_PoPaM made a broader cluster with PANTR, but PANTR was far from all three organisms viz. PONPY, PAPHA, and MACMU compared to distances among the three. On the other hand, 9B showed that EC_2_GPPoPaMH was generated in Type-2 (similar to Figure 7B) as pairwise distances of its components were almost the same. Pairwise distances of PONPY, PAPHA, and MACMU were significantly smaller in Type-1 than Type-2. CANLF was closer to all above-mentioned six organisms in Type-1 than Type-2 (Figure 9).

**Figure 9:**
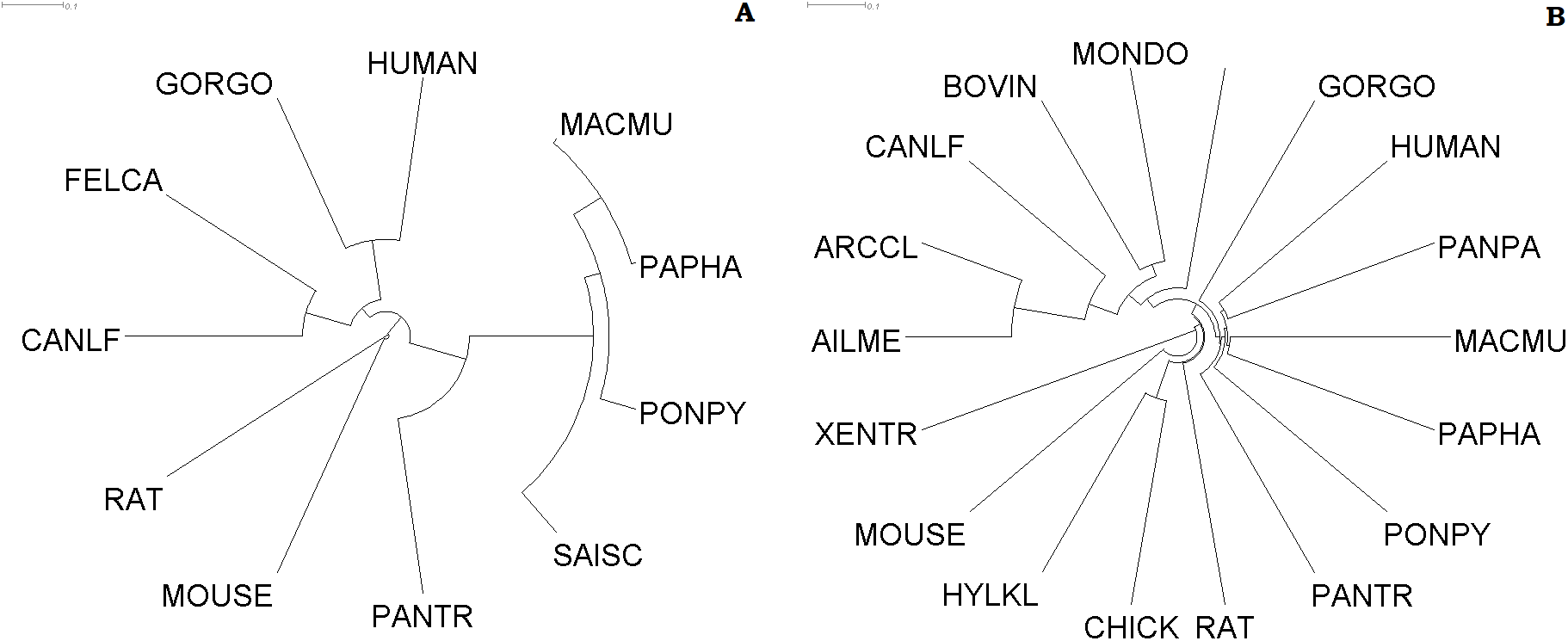
Clustering of organisms based on polar, non-polar amino acid sequence homology of Type-1 (A) and Type-2 (B) taste receptors

### 3.7. Amino acid frequency based correlation analysis, hierarchical relationship, and clustering

*Correlation analysis:* It was found that the highest frequency of amino acids (AAC) in all the taste receptor sequences was obtained in the case of Leucine (L). Based on frequency distribution of twenty amino acids over the Type-1 sequences, out of 380 all possible pairs, 17 pairs of strongly correlated (either positively or negatively, absolute value of correlation coefficient ≥ 0.8) amino acids were found (Table 5). It was noted that amino acids A, R, N, D, C, Q, E, G, H, I, L, K, M, F, P, S, T, W, Y, and V were numbered as 1, 2, 3, … 20, respectively (as plotted in Figure 10).

**Table 5:** Positive or negative strong correlations (absolute value of correlation coefficient: (r) ≥ 0.8) of amino acids based on frequency distribution across the Type-1 taste receptor sequences.

| Sl. No. | Amino acid pairs | Type of strong correlation | Sl. No. | Amino acid pairs | Type of strong correlation |
| --- | --- | --- | --- | --- | --- |
| 1 | (A, N) | Negative | 10 | (N, Y) | **Positive** |
| 2 | (A, G) | **Positive** | 11 | (G, I) | Negative |
| 3 | (A, I) | Negative | 12 | (G, W) | **Positive** |
| 4 | (A, L) | **Positive** | 13 | (G, Y) | Negative |
| 5 | (A, W) | **Positive** | 14 | (I, L) | Negative |
| 6 | (A, Y) | Negative | 15 | (I, Y) | **Positive** |
| 7 | (N, G) | Negative | 16 | (L, T) | Negative |
| 8 | (N, I) | **Positive** | 17 | (L, Y) | Negative |
| 9 | (N, W) | Negative |  |  |  |

**Figure 10:**
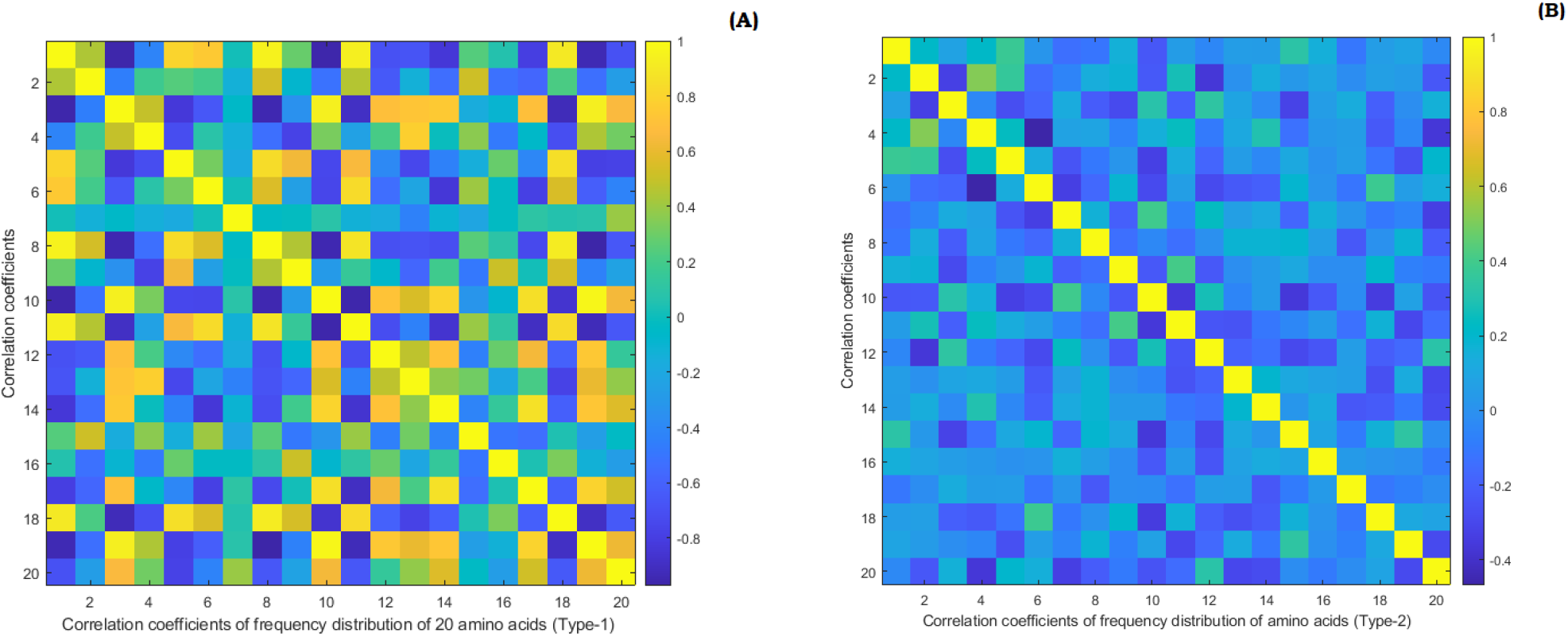
Correlation coefficient matrices of all the Type-1 (A) and Type-2(B) taste receptor sequences. Yellow color represents the Correlation coefficient 1.

On the other hand, no strong correlation (with absolute value of correlation coefficient greater than 0.8) of frequency distribution of amino acids was noticed across the Type-2 taste receptor sequences. It was noted that the highest absolute value of correlation coefficient 0.5167 was obtained for a pair of amino acids (R,D) for the Type-2 taste receptor sequences.

*Hierarchical relationship and clustering:* Three sweet taste receptor members of different organisms were clearly clustered into three broad clusters based on amino acid composition (Figure 11), as it was observed in the cases of sequence homology and polarity based similarity. Hierarchical relationship of Type-2 taste receptor sequences based on amino acid composition is given in the **Supplementary file-6**.

**Figure 11:**
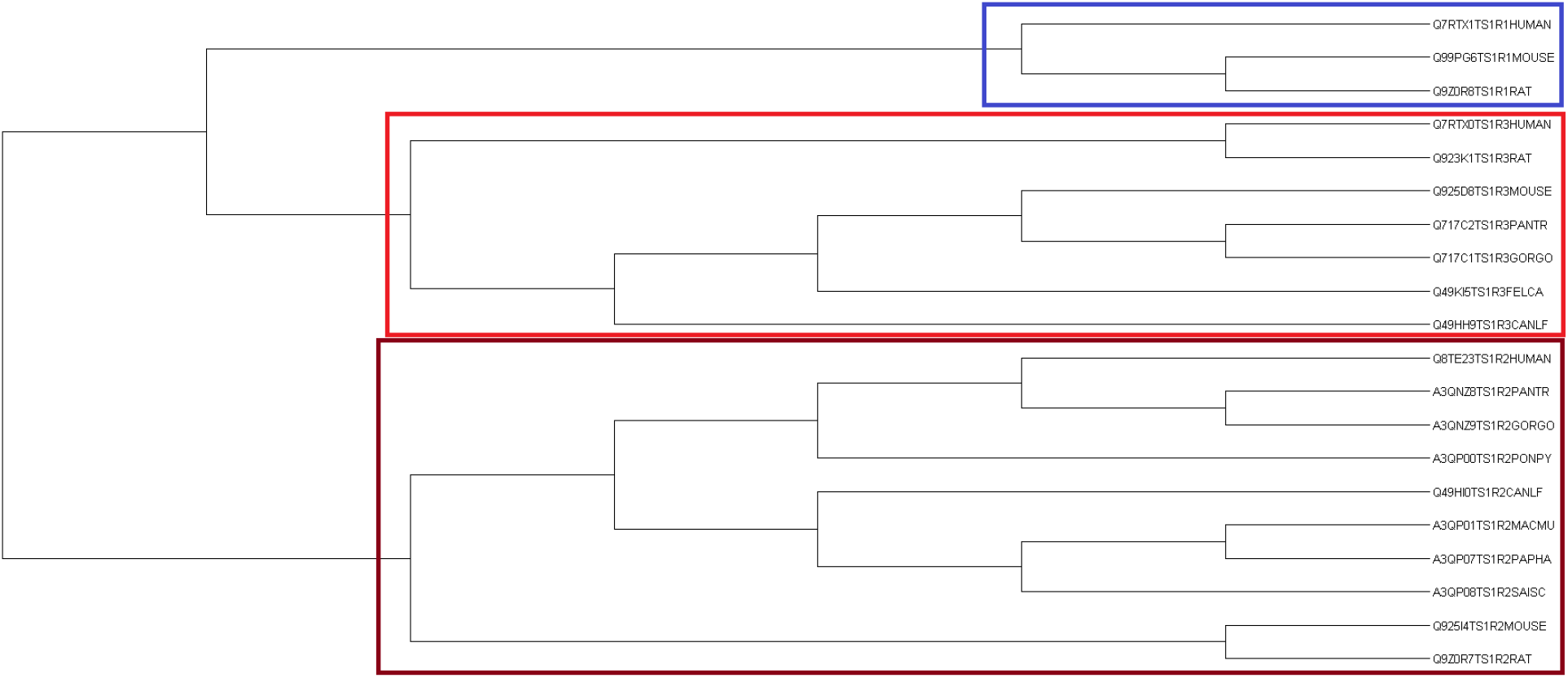
Hierarchical relationship of Type-1 taste receptors based on amino acid frequency and composition

Comparison between Type-1 and Type-2 clusters based on AAC has much similarity with that based on MSA (Figure 7). Here also EC_1_GP and EC_1_PoPaM were formed in Type-1 (Figure 12A) and EC_2_GPPoPaMH in Type-2. Distance between MACMU and PAPHA was very small in Type-1 and both of them were away from EC_1_GP. Comparison of HUMAN and CANLF between Type-1 and Type-2 remained the same as MSA (Figure 12).

**Figure 12:**
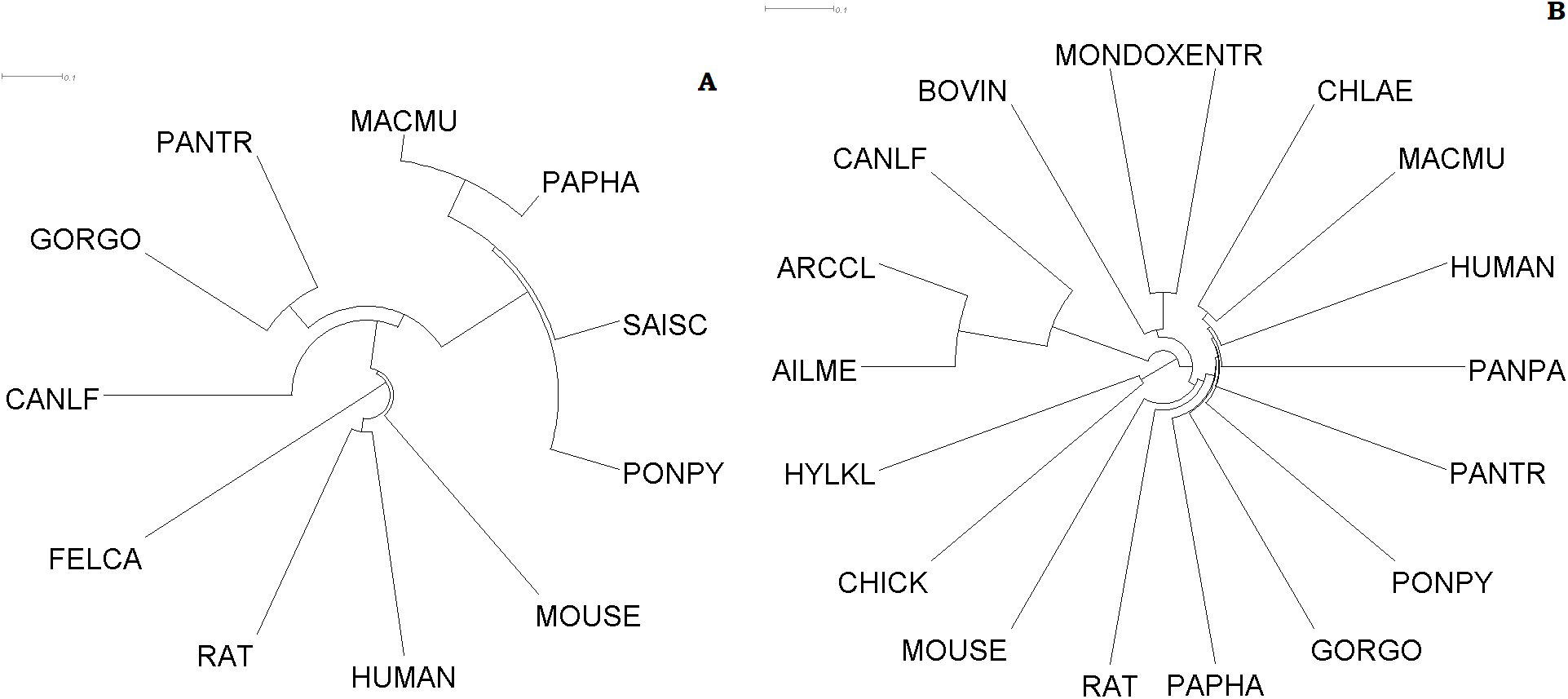
Clustering of organisms based on amino acid frequency distribution Type-1 (A) and Type-2 (B) taste receptor sequences

### 3.8. Structural and physicochemical features based statistical analysis, hierarchical relationship, and clustering

*Significance test:* Out of 1182 PC features, 406 were found to be significantly different between Type-1 and Type-2 from Mann Whitney U test. Among these 406 features, 14, 96, 5, 15, 29, 32, 162, 26, and 27 features were from AAC, DPC, GAAC, GDPC, Moral autocorrelation, CTDC, Conjoint Triad, SoC number, and PAAC.

*Correlation analysis:* These 406 significant features were considered for correlation analysis. Feature-pairs having absolute value of Pearson correlation coefficients greater than 0.8 at significance level of 0.01 were considered to be strongly correlated (Figure 13). 3000 feature-pairs were found to be strongly correlated for Type-1 taste receptors in contrast to 50 feature-pairs in Type-2.

**Figure 13:**
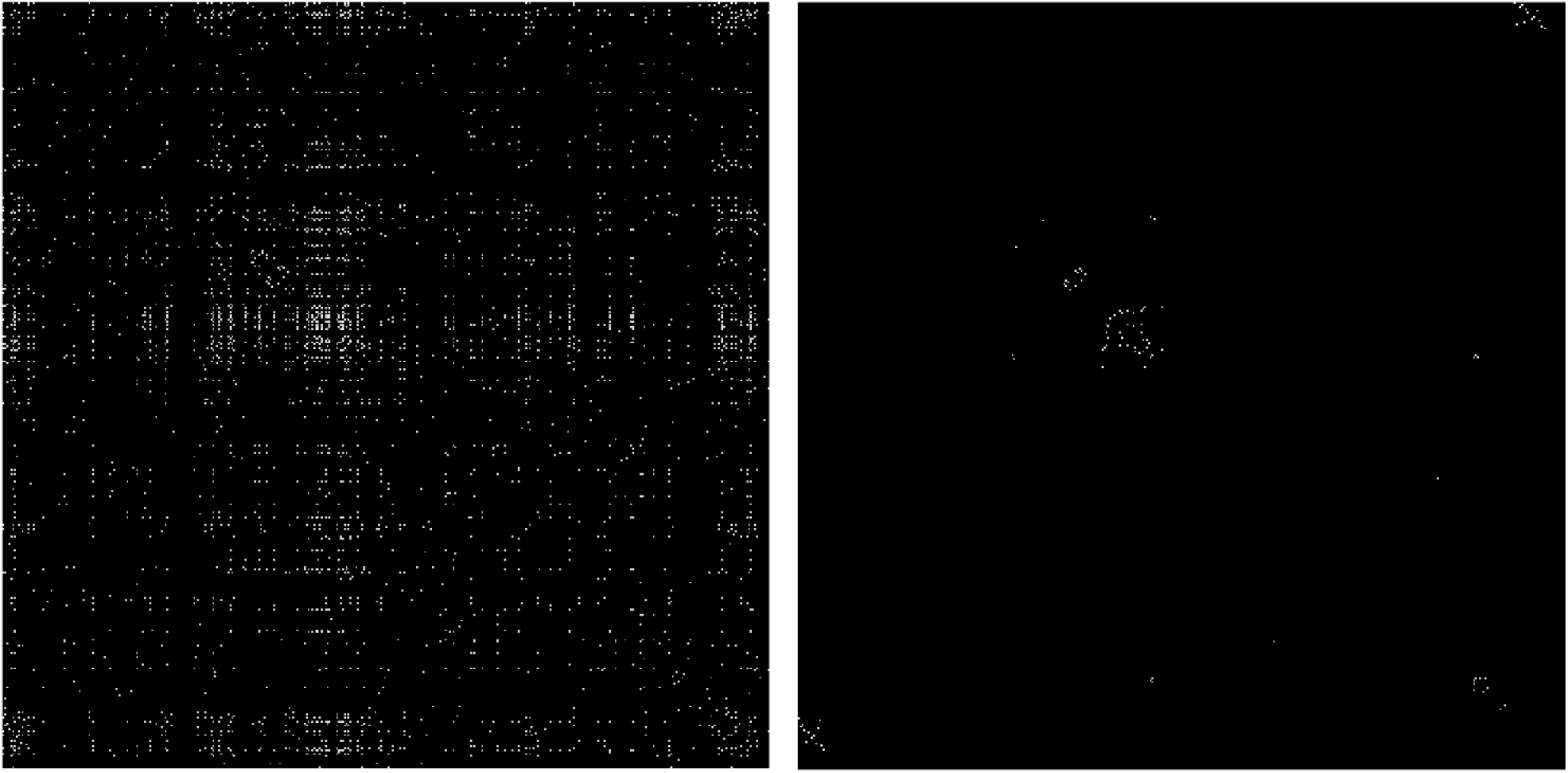
Correlation matrices of size 406 × 406 of Type-1 (left) and Type-2 (right). Figures of correlation matrices are binary, where absolute value of correlation coefficients < 0.8 are denoted by black and ≥ 0.8 are white.

Among the strongly correlated features, 38 feature-pairs were found common in both Type-1 and Type-2 (Table 6). It was followed that two different descriptors viz. amino acid composition and pseudo-amino acid composition were strongly correlated in 12 pairs. Both the features belong to Moran autocorrelation in 6 pairs and CTDC in 17 pairs.

**Table 6:** List of common 38 structural and physicochemical feature-pairs, which were found to be strongly correlated in both the Type-1 and Type-2 taste receptors sequence.

| Common structural and physicochemical feature-pairs (38) | Descriptor paired with |  |
| --- | --- | --- |
| ('A','Xc1.A') | AAC | PAAC |
| ('C','Xc1.C') | AAC | PAAC |
| ('D','Xc1.D') | AAC | PAAC |
| ('E','Xc1.E') | AAC | PAAC |
| ('F','Xc1.F') | AAC | PAAC |
| ('G','Xc1.G') | AAC | PAAC |
| ('K','Xc1.K') | AAC | PAAC |
| ('L','Xc1.L') | AAC | PAAC |
| ('P','Xc1.P') | AAC | PAAC |
| ('Q','Xc1.Q') | AAC | PAAC |
| ('R','Xc1.R') | AAC | PAAC |
| ('S','Xc1.S') | AAC | PAAC |
| ('negativecharge', 'charge.G3') | GAAC | CTDC |
| ('uncharge', 'uncharger.uncharger') | GAAC | GDPC |
| ('CHAM820101.lag1', 'BIGC670101.lag1') | Moran Autocorrelation | Moran Autocorrelation |
| ('CHAM820101.lag6', 'CHOC760101.lag6') | Moran Autocorrelation | Moran Autocorrelation |
| ('CHAM820101.lag6', 'BIGC670101.lag6') | Moran Autocorrelation | Moran Autocorrelation |
| ('CHAM820101.lag9', 'CHOC760101.lag9') | Moran Autocorrelation | Moran Autocorrelation |
| ('CHAM820101.lag23', 'BIGC670101.lag23') | Moran Autocorrelation | Moran Autocorrelation |
| ('CHOC760101.lag6', 'BIGC670101.lag6') | Moran Autocorrelation | Moran Autocorrelation |
| ('hydrophobicity_PRAM900101.G1', 'polarity.G3') | CTDC | CTDC |
| ('hydrophobicity_PRAM900101.G1', 'solventaccess.G2') | CTDC | CTDC |
| ('hydrophobicity_PRAM900101.G2', 'polarity.G2') | CTDC | CTDC |
| ('hydrophobicity_PRAM900101.G3', 'hydrophobicity_PONP930101.G3') | CTDC | CTDC |
| ('hydrophobicity_PRAM900101.G3', 'hydrophobicity_ENGD860101.G3') | CTDC | CTDC |
| ('hydrophobicity_PRAM900101.G3', 'polarity.G1') | CTDC | CTDC |
| ('hydrophobicity_ARGP820101.G3', 'hydrophobicity_ZIMJ680101.G3') | CTDC | CTDC |
| ('hydrophobicity_PONP930101.G3', 'hydrophobicity_ENGD860101.G3') | CTDC | CTDC |
| ('hydrophobicity_PONP930101.G3', 'polarity.G1') | CTDC | CTDC |
| ('hydrophobicity_CASG920101.G1', 'hydrophobicity_FASG890101.G3') | CTDC | CTDC |
| ('hydrophobicity_ENGD860101.G3', 'polarity.G1') | CTDC | CTDC |
| ('normwaalsvolume.G1', 'polarity.G2') | CTDC | CTDC |
| ('normwaalsvolume.G1', 'polarizability.G1') | CTDC | CTDC |
| ('normwaalsvolume.G3', 'polarizability.G3') | CTDC | CTDC |
| ('polarity.G2', 'polarizability.G1') | CTDC | CTDC |
| ('polarity.G3', 'solventaccess.G2') | CTDC | CTDC |
| ('charge.G2', 'Schneider.lag8') | CTDC | SoC number |
| ('g2.g6.g7', 'g6.g7.g2') | Conjoint Triad | Conjoint Triad |

*Hierarchical relationship and clustering:* Similar to previous three hierarchical relationships, three members of Type-1 formed three disjoint clusters (Figure 14). **Supplementary file-7** depicts the hierarchical relationshipns among Type-2 taste receptors based on PC.

**Figure 14:**
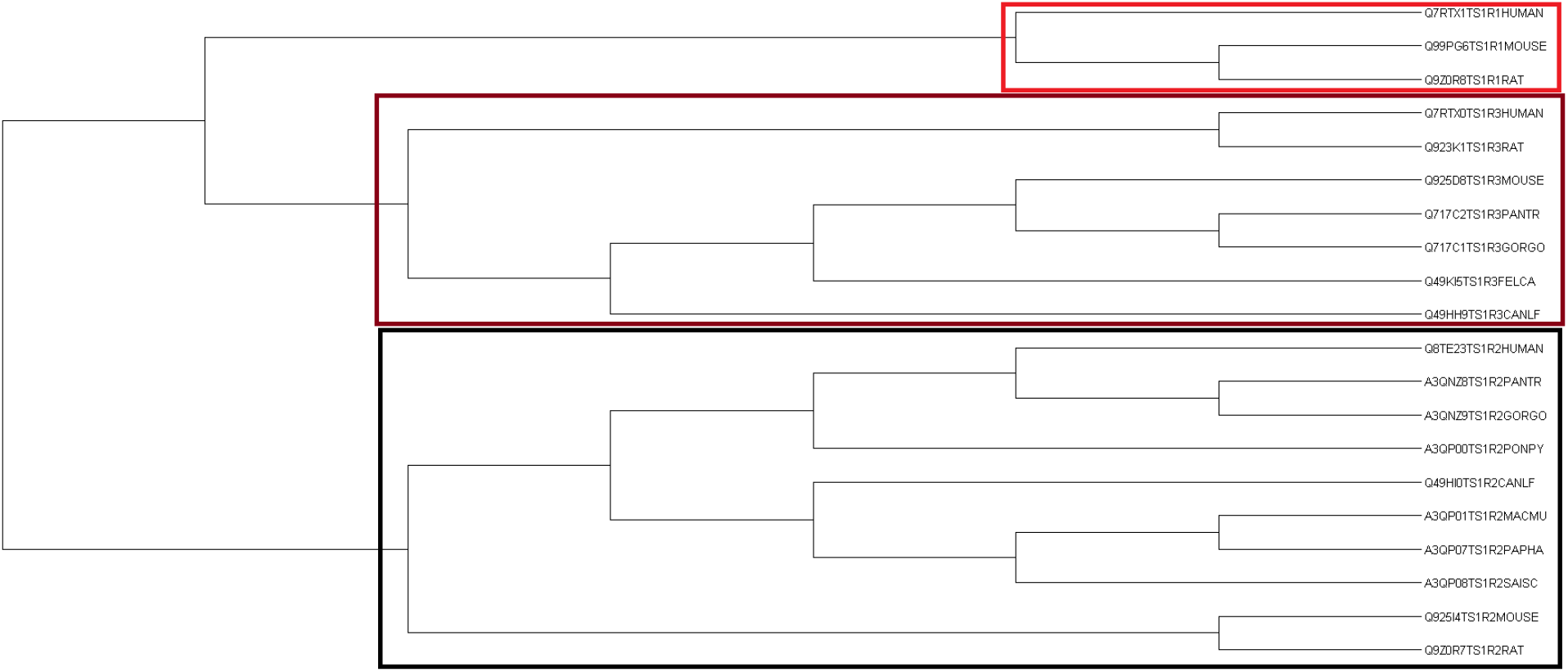
Hierarchical relationship of Type-1 taste receptors based on physicochemical properties

Comparison between Type-1 and Type-2 clusters based on PC is exactly the same as AAC (Figure 12) except for HUMAN and CANLF. From Figure 15 it was followed that HUMAN was far apart from both GORGO and PANTR (or EC_1_GP) as well as EC_1_PoPaM. It can be noted that unlike Figure 12, distances between CANLF and GORGO, PANTR, PONPY, PAPHA, MACMU, and HUMAN in Type-1 were almost similar to Type-2 (Figure 15).

**Figure 15:**
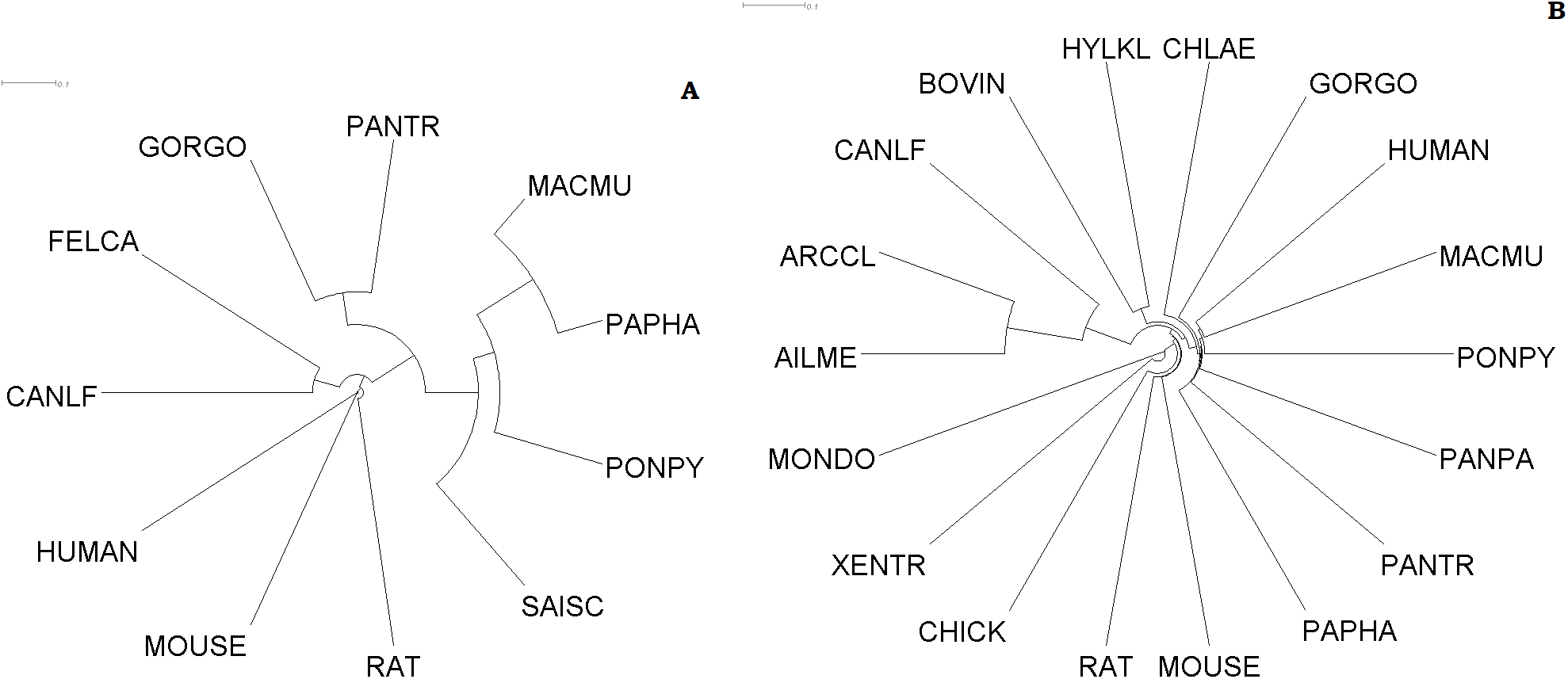
Clustering of organisms based on structural and physicochemical features extracted from Type-1 (A) and Type-2 (B) taste receptor sequences

### 3.9. Biophysical features based statistical analysis, hierarchical relationship, and clustering

*Significance test:* It was found that all the nine biophysical features (BP) were found to be significantly different between Type-1 and Type-2 from Mann Whitney U test.

*Correlation analysis:* For Type-1 taste receptors, only one feature pair (Theoretical PI, total number of negatively charged residues) was found to be strongly correlated with a positive correlation coefficient greater than 0.8. In the case of Type-2 taste receptors, no pair was detected to have strong correlation.

One of the biophysical features viz. Shannon entropy illustrates the disorderliness of amino acid frequency distribution across the taste receptor sequences.

*Amino acid frequency based Shannon entropy:* Amino acid frequency based Shannon entropy (SE) was calculated for all the taste receptor sequences of Type-1 and Type-2. Shannon entropy of the Type-1 taste receptor sequences was ranging from 0.937 to 0.9705, very close to 1. Also, the SE of Type-2 taste receptor proteins were found to be lying in the interval [0.90234, 0.94725].

In both types of taste receptors such a high SE showed a very high level of disordered-frequency distribution of amino acids over the taste receptors. Furthermore, it noticed that the SE of the Type-1 taste receptor proteins viz. Q7RTX0—TS1R3_HUMAN, Q923K1—TS1R3_RAT, Q9Z0R8—TS1R1_RAT, Q99PG6—TS1R1_MOUSE, Q7RTX1—TS1R1_HUMAN, A3QNZ9—TS1R2_GORGO, A3QNZ8—TS1R2_PANTR, A3QP08—TS1R2_SAISC, Q8TE23—TS1R2_HUMAN, A3QP00—TS1R2_PONPY, Q9Z0R7—TS1R2_RAT, Q925I4—TS1R2_MOUSE, A3QP07—TS1R2_PAPHA, A3QP01—TS1R2_MACMU, and Q49HI0—TS1R2_CANLF were higher than the maximum SE obtained for all the 189 Type-2 taste receptors. It was turned out that amino acid frequency distribution was more strongly disordered in Type-1 than that of the Type-2 taste receptor sequences. The highest SEs for Type-1 (0.9705) and Type-2 (0.9472) receptors were noticed in Q49HI0—TS1R2_CANLF and sp—Q645U7—T2R60_PONPY, respectively.

*Hierarchical relationship and clustering:* Unlike the previous four (MSA, PNP, AAC, PC) here three members of Type 1 taste receptors do not form separate clusters (Figure 16). Hierarchical relationship of Type-2 taste receptors is supplied in the **Supplementary file-8**.

**Figure 16:**
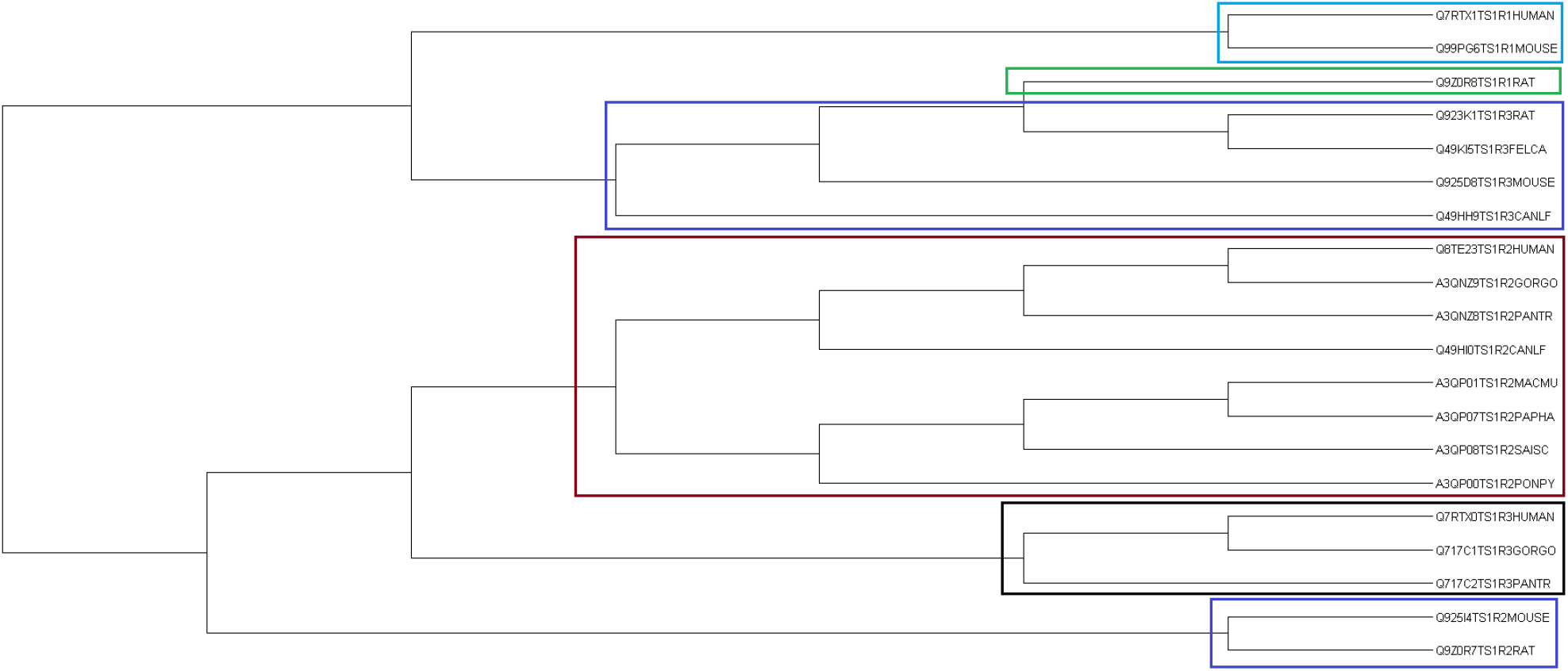
Hierarchical relationship of Type-1 taste receptors based on biophysical properties

HUMAN, GORGO, and PANTR belonged to an extended cluster (EG_1_HGP) in Type-1 (Figure 17). MACMU and PAPHA were paired as they were very close to each other. Here also EC_1_PoPaM (extended cluster of MACMU, PAPHA, and PONPY) was present and the same was away from ECLHGP. EC_2_GPPoPaMH was formed in Type-2 similar to MSA (Figure 7), PNP (Figure 9), AAC (Figure 12), and PC (Figure 15). Unlike MSA (Figure 7), PNP (Figure 9), and AAC (Figure 12), CANLF was closer to all above-mentioned six organisms in Type-2 than Type-1.

**Figure 17:**
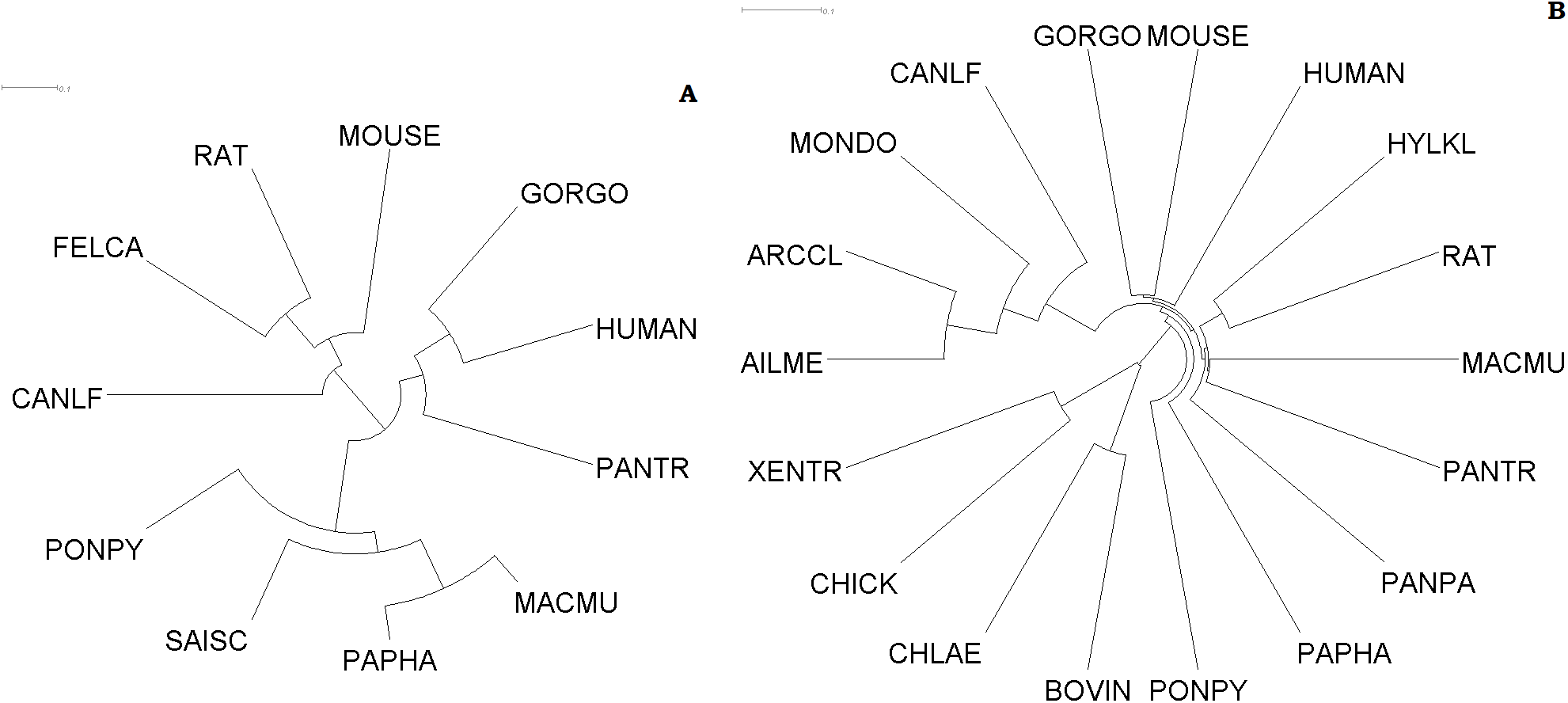
Clustering of organisms based on biophysical features extracted from the Type-1 (A) and Type-2 (B) taste receptors

### 3.10. Nearness of organisms based on taste receptors’ features

#### 3.10.1. Cumulative clustering of organisms (Type-1)

Out of five individual clusters based on SNP (Figure 4A), MSA (Figure 7A), PNP (Figure 9A), PC (Figure 15A), and BP (Figure 17A), FELCA was paired with CANLF in three clusters viz. MSA, PNP, and PC. From the cluster of the SNP, it seems that FELCA and CANLF were far apart. But, the distance between FELCA and CANLF was actually lowest in SNP among all five and highest in PC. Furthermore, FELCA and CANLF were significantly closer to HUMAN, GORGO and PANTR compared to MACMU, PAPHA, and PONPY in SNP. In cumulative clustering (Figure 18A), FELCA and CANLF formed a distinct cluster, which was adjacent to the cluster containing HUMAN, GORGO, and PANTR.

**Figure 18:**
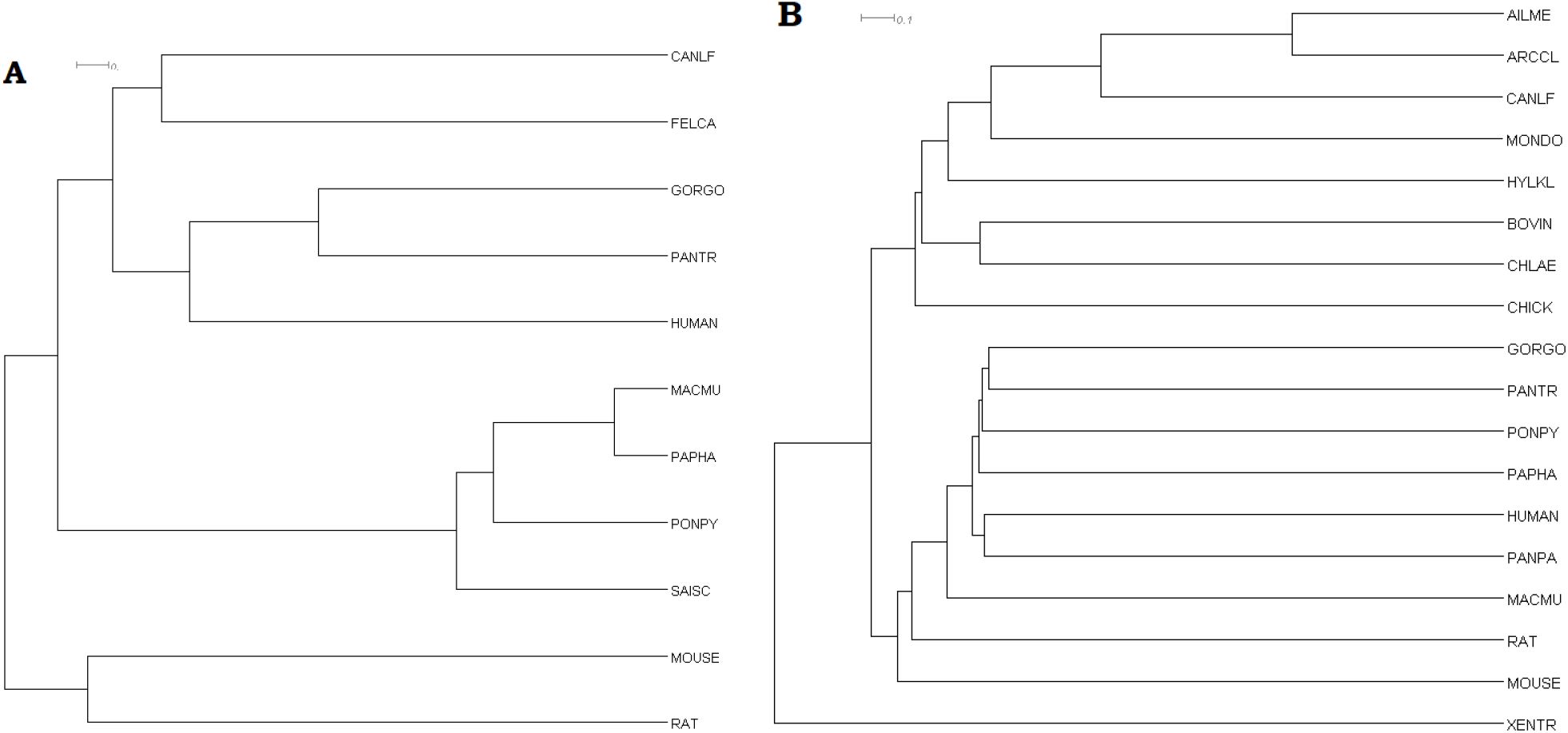
Cumulative clustering of organisms based on the features extracted from the Type-1 (A), and Type-2 (B) taste receptors

GORGO and PANTR were paired in PC and MSA while HUMAN, GORGO, and PANTR formed an extended cluster in BP and SNP. Pairwise distances among HUMAN, GORGO, and PANTR were significantly less in SNP than other four figures. As HUMAN was markedly away from PANTR in PC, MSA, and PNP as well as HUMAN was distant from GORGO in PC, Figure 18A shows GORGO and PANTR created a distinct cluster with HUMAN next to it.

Each of the five clusters had an extended cluster which contained only MACMU, PAPHA, PONPY and SAISC. MACMU and PAPHA especially were very close in all figures. Further, the clustering hierarchy of these four organisms were identical in four clusters except BP (Figure 17A). In BP, SAISC became close to the pair MACMU-PAPHA instead of PONPY. Eventually these four organisms were also grouped together in Figure 18A and their hierarchy was replicated as in four figures (MSA, SNP, PNP, and PC).

MOUSE and RAT formed a distinct cluster only in SNP where the distance between them was markedly less than other four clusters (MSA, PNP, PC, and BP). MOUSE was adjacent to the smallest cluster containing RAT in BP. In the rest of the three figures (MSA, PNP, and PC), MOUSE and RAT were added individually after formation of the extended cluster containing eight organisms of Type-1 viz. CANLF, FELCA, GORGO, PANTR, MACMU, PAPHA, PONPY, and SAISC. Finally, MOUSE and RAT formed a distinct cluster in Figure 18A and this pair was away from all other organisms. Inter group comparison of five individual clusters revealed MSA and PC were very much similar in Type-1.

#### 3.10.2. Cumulative clustering of organisms (Type-2)

AILME and ARCCL made a distinct cluster in all of the five figures viz. MSA (Figure 7B), SNP (Figure 4B), PNP (Figure 9A), PC (Figure 15A), and BP (Figure 17B). CANLF was adjacent to the pair AILME-ARCCL and generated an extended cluster in four figures viz. SNP, MSA, PNP, and PC. In BP, MONDO was adjacent to the pair AILME-ARCCL and CANLF was next to the extended cluster containing AILME, ARCCL, and MONDO. AILME, ARCCL, CANLF, MONDO, and HYLKL formed an extended cluster (EC_2_AiArCaMoHy) with very small pairwise distances among them in SNP. Furthermore, EC_2_AiArCaMoHy was away from other clusters in SNP. This leads to formation of EC_2_AiArCaMoHy in the cumulative clustering of Type-2 (Figure 18B).

BOVIN and CHLAE made a distinct cluster in BP and SNP. BOVIN, CHLAE, and CHICK were clubbed in SNP with very small pairwise distances among them. Extended cluster consisting of BOVIN, CHLAE, and CHICK were adjacent to EC_2_AiArCaMoHy in SNP. This nature of clustering was also reflected in (Figure 18B).

GORGO, PANTR, PONPY, PAPHA, HUMAN, and PANPA being almost equidistant to each other, formed an extended cluster in Type-2 (EC_2_GPPoPaHPan) in all five clusters. Specifically, HUMAN and PANPA made a distinct cluster in SNP and PNP. Negligible pairwise distances among GORGO, PANTR, PONPY, and PAPHA in SNP made them develop an extended cluster in (Figure 18B), while HUMAN was paired with PANPA. The pair HUMAN-PANPA was adjacent to clusters containing GORGO, PANTR, PONPY, PAPHA in Figure 18B. MACMU was slightly away from the cluster of GORGO, PANTR, PONPY, PAPHA as well as the pair HUMAN-PANPA in Figure 18B because of MACMU’s position in SNP. Although MOUSE and RAT formed a distinct cluster in SNP, considering all five figures they did not form any pair in Figure 18B.

XENTR was away from all extended clusters in Figure 18B as XENTR was added after all the remaining seventeen organisms of Type-2 were grouped in SNP, MSA, and PNP. Even in PC, XENTR was added after sixteen organisms of Type-2 were grouped.

#### 3.10.3. Cumulative clustering of common organisms (Type-1 and Type-2)

Nine common organisms viz. HUMAN, MACMU, PAPHA, PONPY, MOUSE, RAT, CANLF, GORGO, and PANTR were clustered into four clusters (Figure 19).

**Figure 19:**
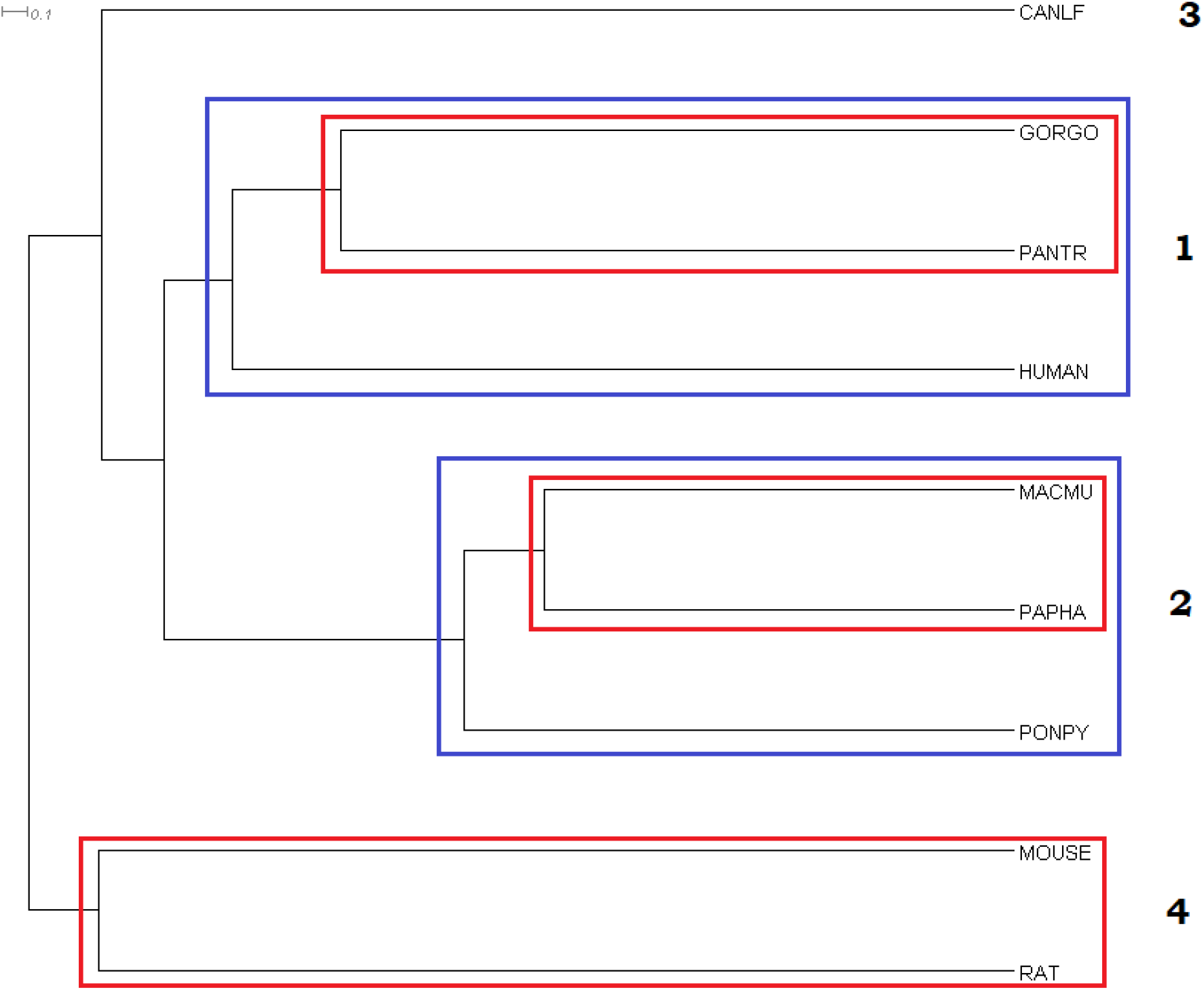
Clustering of common organisms associated to both Type-1 and Type-2 receptors

It was observed from (Figure 19) that three clusters viz. EC_HGP (extended cluster of HUMAN, GORGO, PANTR), ECP_PoPaM (extended cluster of PONPY, PAPHA, and MACMU) and the pair RAT-MOUSE were turned out to be the same as obtained in the cumulative clustering for Type-1 (Figure 18A), although adjacency between the first two clusters (EC_HGP and EC_PoPaM) slightly changed in (Figure 19). EC_HGP was next to CANLF in (Figure 18A), while EC_HGP was adjacent to EC_PoPaM in (Figure 19). It is noteworthy that GORGO, PANTR were paired in EC_HGP and PAPHA, MACMU were paired in EC_PoPaM of Figure 18A, which were mirrored in Figure 20. On the other hand, EC_HGP and EC_PoPaM did not exist separately in cumulative clustering of Type-2 (Figure 18B), instead all the six organisms viz. HUMAN, GORGO, PANTR, PONPY, PAPHA, and MACMU made an extended cluster (EC_HGPPoPaM). The change of adjacency between EC_HGP and EC_PoPaM in Figure 19 from Figure 18A might arise from the fact that EC_HGPPoPaM was distant from CANLF in (Figure 18B). In both Figure 18A and Figure 19, the pair MOUSE and RAT were added after all other organisms were grouped.

**Figure 20:**
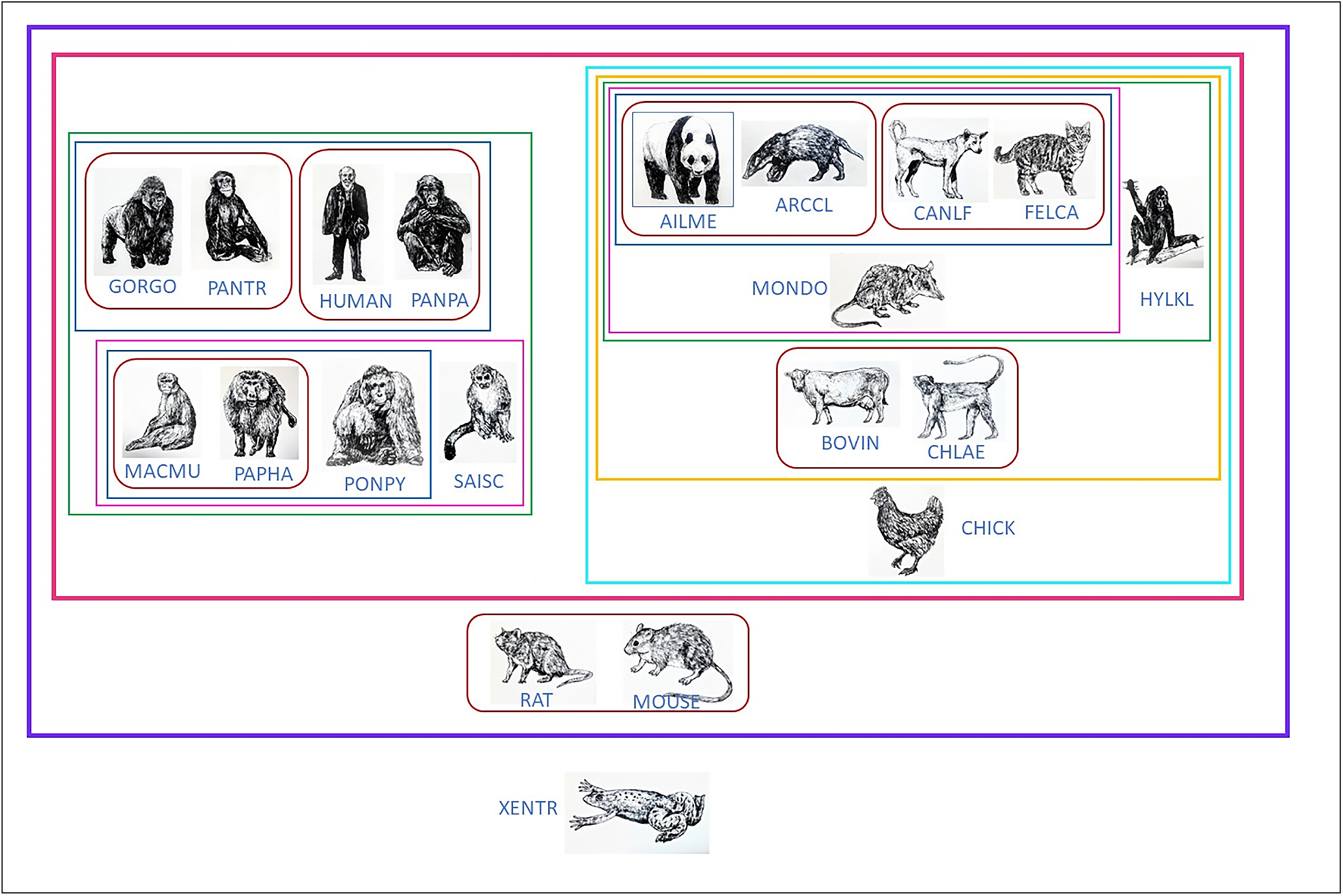
Possible functional proximity of 20 organisms based on taste receptors’ features

Above discussion clearly indicates the importance of both Type-1 and Type-2 sequences in obtaining the proximity among the organisms.

#### 3.10.4. Possible functional proximity of organisms

Combining Figure 18A, Figure 18B, and Figure 19 formed following pairs {HUMAN, PANPA}, {GORGO, PANTR}, {MACMU, PAPHA}, {AILME, ARCCL}, {CANLF, FELCA}, and {BOVIN, CHLAE} (Figure 20). Two members of each pair shared a very strong nearness. The pair {MOUSE, RAT} was added after all other organisms except XENTR were clubbed (Figure 20). Divergence of Type-2 taste receptor sequences of XENTR made it away from all the rest 19 organisms (Figure 20).

## 4. Conclusions

20 sweet (Type-1) and 189 bitter (Type-2) taste receptors were considered from 20 different vertebrate organisms to comprehend the proximity among the organisms based on similarity and dissimilarity of taste receptor sequences. It was found that the binary representation of polar-nonpolar sequences of all the organisms deviated from random binary sequences. Thus, depicting an ‘‘organized” spatial distribution of polar-nonpolar amino acids along the taste receptor sequences. Furthermore, for both receptor types, the largest variability was observed within the disordered or flexible regions of these proteins as noticed in protein intrinsic disorder disposition. Three members of sweet taste receptors formed three disjoint clusters in all five quantitative features except biophysical feature. Disparity between sweet and bitter taste was reflected by Shannon variability, correlation and significance analysis of biophysical and physicochemcial features. Furthermore, divergence among bitter taste receptor sequences was clearly noted through single nucleotide polymorphism mutations, amino acid homology, polar-nonpolar homology, and amino acid frequency distribution.

Unification of diversification in sweet and bitter taste receptor sequences lead to a classification among 20 organisms. In this study, FELCA and SAISC do not have bitter taste receptors, while nine organisms viz. AILME, ARCCL, HYLKL, MONDO, BOVIN, CHLAE, CHICK, PANPA, and XENTR do not have sweet taste receptors. Still all the aforementioned 11 organisms were hypothesised to be in close proximity to the organisms with which they formed clusters in sweet and bitter taste, individually. Out of 18 mammals, 17 belong to subclass Eutheria and one (MONDO) belong to subclass Metatheria. Among 17 Eutherian mammals 10 are Primate, 4 Carnivora (CANLF, FELCA, AILME, ARCCL), 2 Rodentia (RAT and MOUSE), and 1 Artiodactyla (BOVIN). Among 10 primates, 8 of them viz. GORGO, PANTR, HUMAN, PANPA, MACMU, PAPHA, PONPY, and SAISC organisms were grouped together. Four carnivora were clubbed together. Although MONDO belong to a different subclass (order: Didelphimorphia), still it made an extended cluster with order Carnivora. Furthermore, BOVIN and CHALE were paired based on nearness of taste receptors, though BOVIN and CHLAE belonged to two different orders under subclass Eutheria. CHICK (class: Aves) was adjacent to a broad cluster containing 4 Carnivora, 2 Primate, 1 Didelphimorphia, and 1 Artiodactyla in concordance with taste receptors inspite of them being hierarchically distant. Rodent organisms viz. RAT and MOUSE placed together outside the extended cluster comprising of two disjoint clusters. Altogether distinct characteristics of bitter taste receptors of the amphibian organism, XENTR made it away from the rest of the clusters. This study encompasses the nearness of varied organisms, thus, portraying a customary standard of proximity among organism based on taste receptors. This study further hypothesised that ligand binding affinity of sweet/bitter taste molecules in the taste receptors of any proximal pair of organisms (such as HUMAN and PANPA) would be similar.

## Acknowledgements

Authors are very grateful to the laboratory supporters Mr. Ashim Dhar and Mrs. Piyali Maitra for their logistic assistance. Authors would like to thank the Department of Biotechnology, Govt. of India (BT/PR26647/NNT/28/1365/2017), Indian Space Research Organization (ISRO/RES/3/827/19-20), and Indian Statistical Institute (ISI) (ISI/TAC/PROJECT 1/ 2020-21) for their financial support.

## Author contributions statement

AG and SSH conceived the problem and theoretical experiments. SSH, VNU, MS, SC, and DN executed the results and performed the analysis. DN and PB performed statistical analyses. SSH, DN, VNU, MS, and SC wrote the initial draft. All authors reviewed and edited the manuscript. AG supervised the entire project. All the authors checked, reviewed, and approved the final version of the manuscript.

