## Supplementary figures and images for "Possible Functional Proximity of Various Organisms Based on Taste Receptors Genomics"

### Supplementary file-4.png

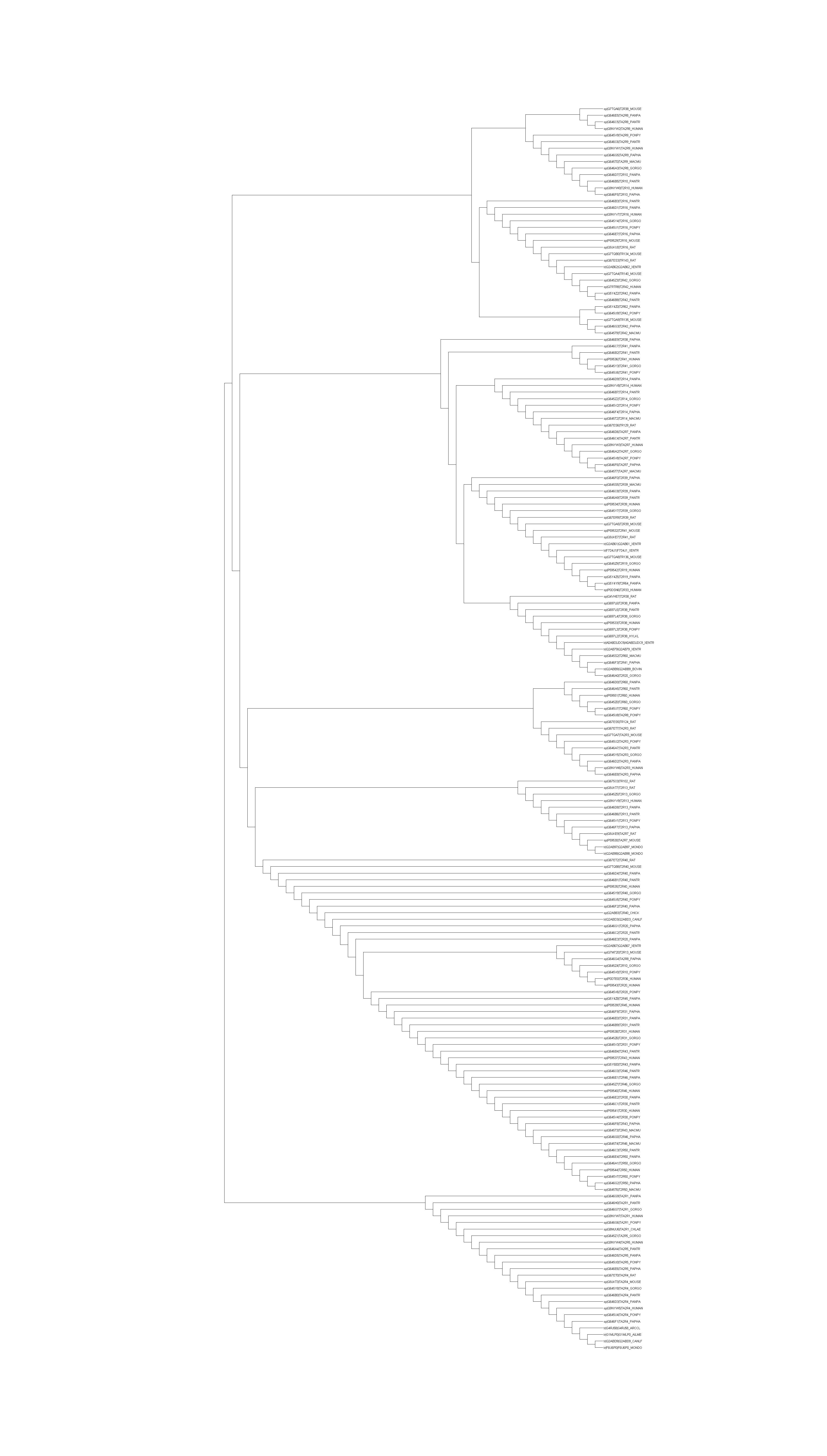

### Supplementary file-5.png

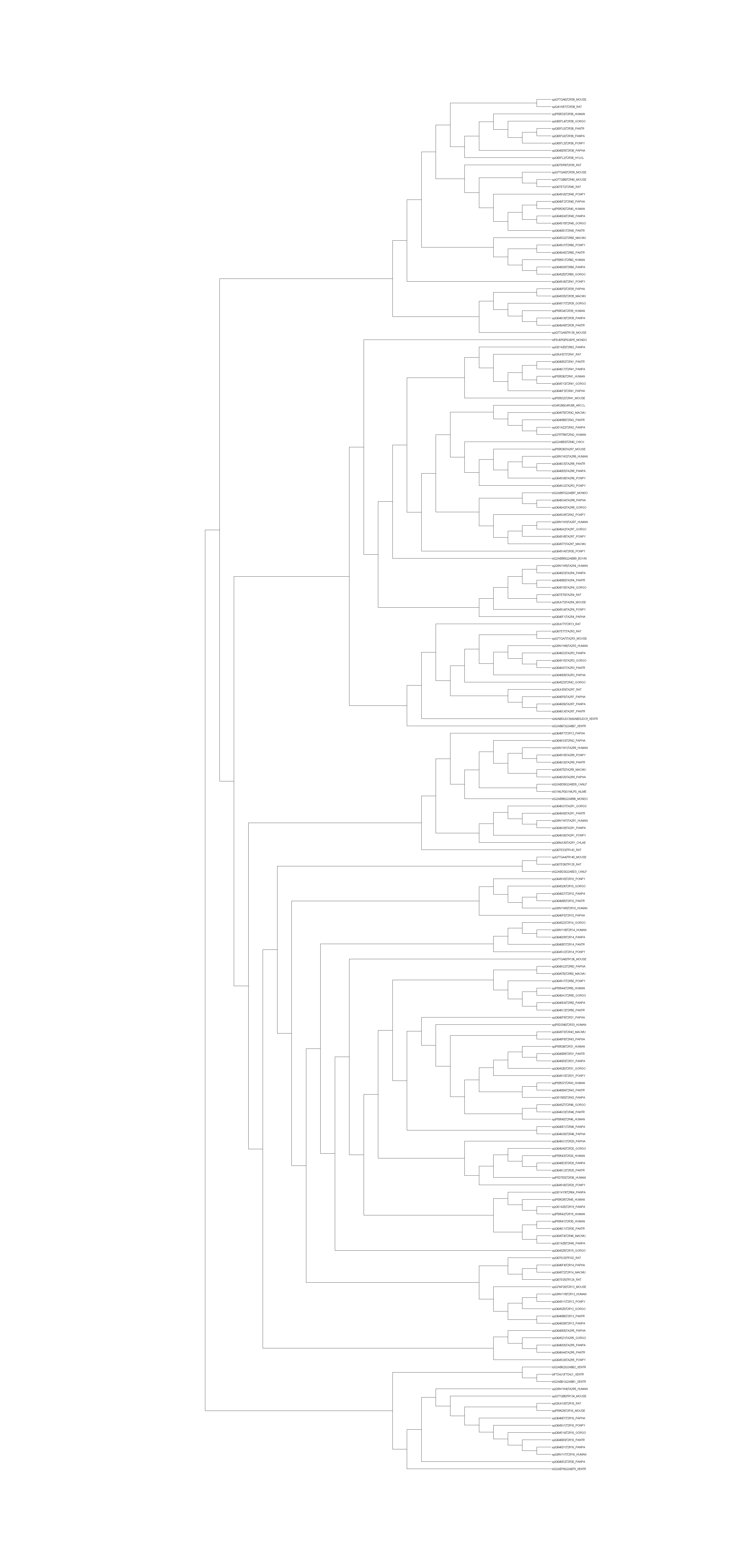

### Supplementary file-6.png

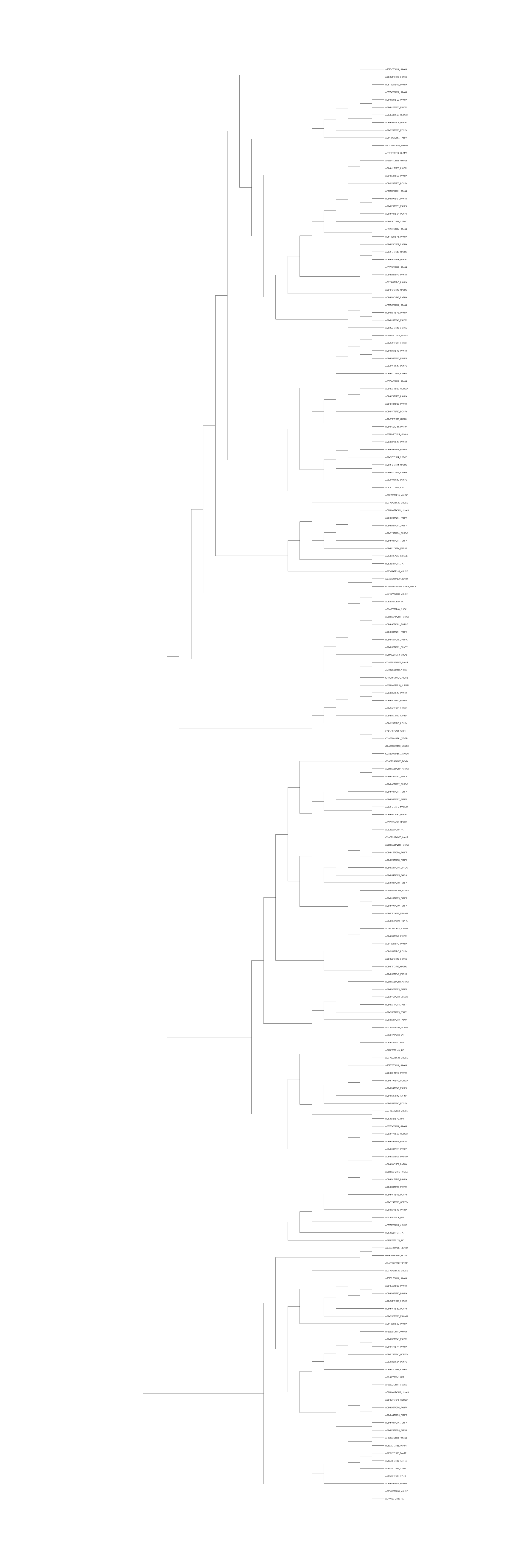

### Supplementary file-7.png

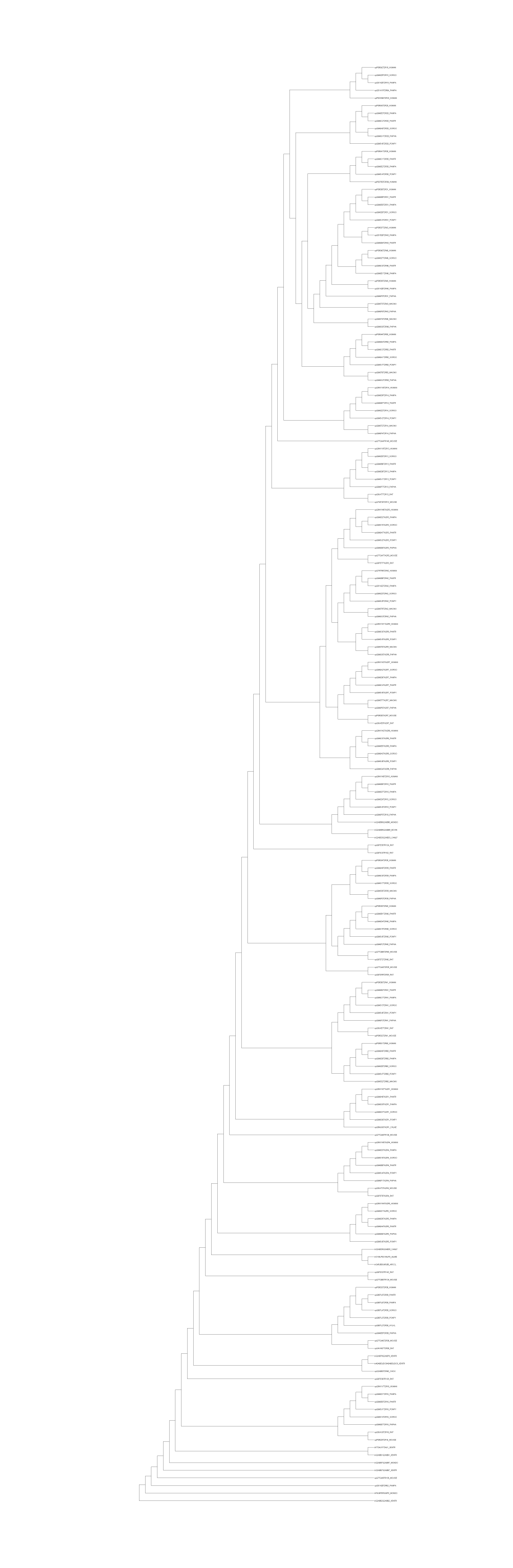

### Supplementary file-8.png

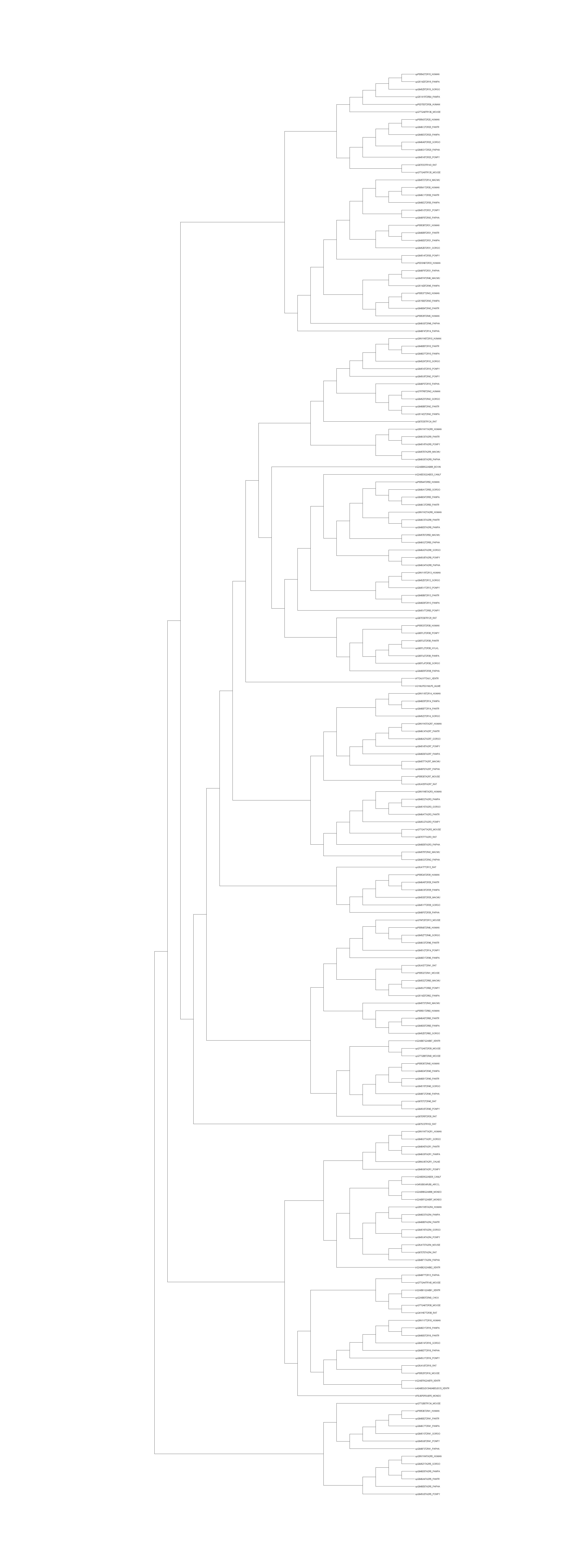
